# Genetic regulation of fasting-induced longevity effects

**DOI:** 10.1101/2025.09.02.673812

**Authors:** Alison Luciano, Laura Robinson, William H. Schott, A. Phillip West, Ron Korstanje, Gary A. Churchill

## Abstract

Research methods for the investigation of the biology of aging have often implicitly generalized strain-specific results. Dietary interventions, such as caloric restriction and periodic fasting, have been shown to enhance metabolic health and extend lifespan in preclinical models. However, inter-individual variation in physiological responses to these interventions, which affects their safety and efficacy when translated to humans, remains poorly understood despite being observed in multiple studies. In this study, we implemented intermittent fasting (IF) for two days per week in 10 inbred strains (n = 800 mice) from the Collaborative Cross (CC). The CC is a multiparent recombinant inbred strain panel that offers a diverse collection of reproducible models to study the genetic control of heterogeneous intervention responses. We conducted longitudinal phenotyping to characterize hundreds of traits, including lifespan, in the CC mice. We demonstrate that sex and genetic background induce variable responses to intermittent fasting across multiple physiological outcomes, including metabolic, hematologic, and immunologic health. Effects of IF on lifespan were sex-specific and variable across genetic backgrounds. Thus we establish that response to IF is genetically determined in an animal model with physiological features similar to humans. We compared our findings in the CC with those from a parallel study of Diversity Outbred (DO) mice, highlighting common predictors of health and lifespan, as well as key differences between the genetically diverse inbred and outbred models. These findings underscore the importance of genetic factors in dietary intervention responses, offering valuable insights for translating intermittent fasting benefits to human health and longevity. Keywords: multiparental populations; gene-by-treatment interaction

## Introduction

Intermittent fasting (IF) is a dietary regimen that cycles between ad libitum (AL) feeding and fasting. Variants of IF include time-restricted feeding (TRF), alternate day fasting (ADF), and periodic fasting (PF) [1]. Although the effects of IF on health and longevity have not been studied as intensively as those of caloric restriction (CR), accumulating evidence points to geroprotective effects [2–9].

Research methods for the investigation of the complex biology of aging have often implicitly generalized strain-specific results. However, genetic modulation of dietary effects on lifespan has been repeatedly observed [10–21], including in our large longitudinal lifespan study of five dietary regimens in outbred mice (Dietary Restriction in Diversity Outbred mice [DRiDO]; Di Francesco *et al.* 2024). Given compelling evidence that genetic background may interact synergistically or antagonistically with dietary restriction, single strain studies of dietary effects are less likely to accurately reflect effects of similar interventions in humans.

Genetic variation in response to dietary intervention is likely attributable to large numbers of loci with small effects. Here we present the Collaborative Cross Longitudinal Study, which was conducted in parallel with DRiDO to demonstrate genetic modification of dietary response in a panel of recombinant inbred strains. We applied the same dietary intervention (IF) to 10 inbred strains from the Collaborative Cross (CC) [22]. The CC population is a multiparental recombinant inbred strain panel created through interbreeding of the genomes of eight founder strains to create independent breeding lines. The finished inbred strains of the CC panel are genetically diverse and reproducible. Extensive longitudinal phenotyping of 800 CC mice of both sexes enabled us to carry out a multisystem exploration of genetic dependence in dietary treatment response across the lifespan. We compared our findings from the CC inbred mouse to the parallel study of dietary restriction in Diversity Outbred (DO) mice, which included a 2-day IF intervention arm. The DO mice are an outbred heterogeneous stock derived from the set of founder strains as the CC.

## Results

Eight hundred mice were randomized to one of two experimental arms (ad libitum feeding [AL, control cohort; n=400] or 2-day intermittent fasting [IF, treatment cohort; n=400]). Mice were weighed weekly and followed with extensive phenotyping until natural death or extreme morbidity. Mice were evenly distributed across 10 Collaborative Cross strains (see Methods) and both sexes. At 6 months of age, with 767 surviving mice, we initiated IF for the treatment cohort and we maintained mice on IF or AL feeding for the duration of their natural lifespan. The experimental design leveraged sex and diverse genetic background to study variable responses to IF across hundreds of physiological outcomes collected longitudinally, including metabolic, hematologic, and immunologic health phenotypes.

### Lifespan response to IF is sexually dimorphic

We examined the effects of IF and AL on lifespan extension by sex, aggregating across the 10 inbred strains. Median lifespan was approximately 24 months in males and 22 months in females. Male mice tended to live longer than female mice in both treatment groups (p=0.038 and p=1.8e-07 in AL and IF, respectively). We observed a significant lifespan effect in response to 2-day intermittent fasting over ad libitum feeding among males but not females (**Fig. 1a**, **Fig. 1b**). Analyses pooling data from both sexes yielded comparable results among males (p=0.00024) and females (p=0.56), as did analyses that accounted for the possibility of intra-cage correlations (male p=0.00075, female p=0.78). The lifespan extension observed in males on IF was modest (median lifespan difference=1.7 months) (**Supplementary Table S1a**). Maximum lifespan, estimated as the 90th percentile, was 30 months in the total sample, 31.1 months among males and 28.8 months among females. Evidence of substantial maximum lifespan extension (*>*1 mo.) was not observed for either sex (**Supplementary Table S1b**). In sensitivity analyses, we re-estimated absolute survival difference estimates of male-driven diet effects by applying restricted mean survival time (RMST) analysis (**Supplementary Table S1c**). The RMST difference in males was 2.02 months (95% CI: 0.76 to 3.28, p=0.002). In contrast, females showed no significant difference in survival between diets (RMST difference: 0.33 months, 95% CI: -0.94 to 1.61, p=0.607), reinforcing the conclusion that IF conferred a sex-specific survival advantage in male CC mice.

**Figure 1:**
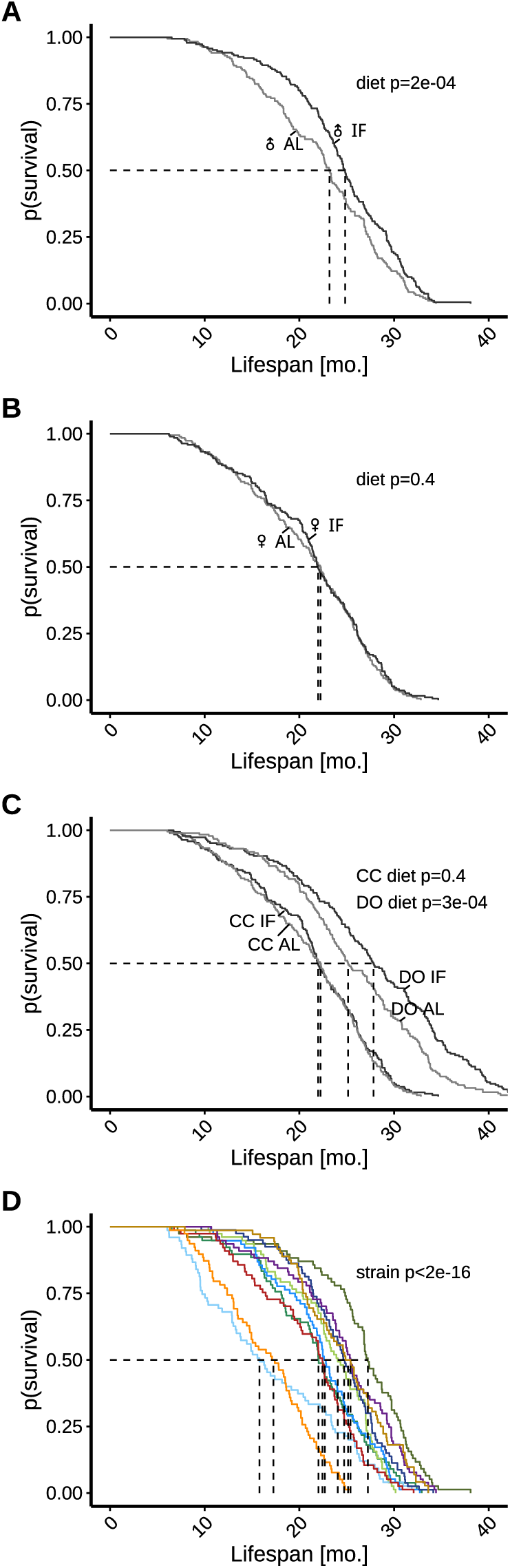
Aggregate lifespan data demonstrates sexually dimorphic response to intermittent fasting (IF) while disaggregated data by genetic background reveals inter-strain variation in IF response. A-B) Study data indicate a significant lifespan extension in males subjected to 2-day intermittent fasting compared to ad libitum feeding, whereas females showed no significant response. C) Female inbred mice exhibit shorter lifespans and weaker IF responses compared to female outbred mice. Outbred data collected from the DRiDO study, which featured high genetic diversity. D) Median survival varied across the 10 inbred strains under control conditions, ranging from 14.1 months in 006/TauUncJ to 26.8 months in 003/UncJ, underscoring the importance of genetic factors in longevity. Abbreviations: CC = Collaborative Cross, DO = Diversity Outbred, mo. = months

### Lifespan response to IF differs in inbred and outbred genetically diverse mice

We previously reported survival data from the DRiDO study for female Diversity Outbred (DO) mice including AL and 2-day IF cohorts (DiFrancesco et al., 2024). The DO are derived from the same founders as the CC strains and thus share the same genetic variants. In addition, the two studies were carried out concurrently in the same mouse facilities. For DRiDO study methods, including information about included animals, the IF dietary intervention, and phenotyping protocols, see [9]. We compared survival times between the two studies and observed that survival times of DO mice were significantly greater than the corresponding CC cohorts (p=1e-13 for AL, p*<*2e-16 for IF). This may reflect an overall fitness cost for inbreeding in the CC mice. We further noted that while IF had no effect on lifespan of female CC mice (p=0.39), there was a significant effect of 2-day IF on lifespan of the female DO mice (p=2.6e-4) (**Fig. 1c**).

### Lifespan and lifespan response to IF varies across CC strains

We observed significant strain-to-strain variation in lifespan of CC mice (p*<*2e-16; **Fig. 1d**). Under AL feeding, median lifespan ranged from 14.1 months in 006/TauUncJ to 26.8 months in 003/UncJ, and sex-specific median lifespan ranged from 17.4 to 27.8 months in males and 13.3 to 26.1 months in females. Under IF feeding, median lifespan ranged from 16.9 months in 006/TauUncJ to 28.7 months in 003/UncJ, and sex-specific median lifespan ranged from 20 to 30.2 months in males and 15.2 to 28.7 months in females (**Supplementary Table S2**). Pairwise tests showed quantitative evidence of heterogeneity in survival by strain (**Supplementary Table S3**).

Study data also support the importance of genetics in lifespan response to IF. Within-strain, Kaplan-Meier curves and interaction contrasts for lifespan response to diet by sex revealed substantial inter-strain variation in IF response (**Supplementary Fig. S1a**). Among females, five strains exhibited a reduced hazard of death on IF, while five strains showed an increased hazard, indicating both beneficial and detrimental effects depending on genetic background. Among males, results were more consistent across strain; eight strains demonstrated reduced hazard of death on IF and only two strains showed increased hazards (**Supplementary Table S4**). Of these comparisons, statistically significant treatment effects were found only among males and all demonstrated lifespan extension on IF (diet p=9e-4, 0.026, and 0.009 for males in 004/TauUncJ, 005/TauUncJ, and 040/TauUncJ, respectively). Results were similar for median lifespan extension or reduction on IF (**Supplementary Fig. S1b**). Strain-specific lifespan variability defined by coefficients of variation (CV; ratio of standard deviation to the mean or sd/m) was similar between AL and IF groups, with standard deviation around a quarter of the mean (**Supplementary Table S5**). Wide intra-strain variation limited power to detect strain specific IF response.

### Heritability of lifespan in CC mice

It is common in preclinical studies to test intervention effects on a single inbred mouse strain, typically C57BL/6. The current study as well as the parallel DRiDO study directly incorporate genetic diversity as feature of the study design. This approach allows for disaggregation of genotypic effects from diet effects via genetic heritability analysis. For mice that lived to at least 6 months of age, genetic background explained 24.74% of variation in lifespan (broad-sense heritability [H2]=0.25, 95% bootstrap CI [0.02, 0.42]), while diet and sex explained only 3.95% of variation. We obtained genotype data and looked at the combined effects of diet, sex and genetic relatedness as estimated by a kinship matrix. Additive genetic effects explained 26.01% of variation in lifespan (narrow-sense heritability [h2]=0.26, 95% bootstrap CI [0.02, 0.45]).

### Genetic background shapes physiological response to intermittent fasting

Longitudinal phenotypic data collected across the lifespan were analyzed for indications that genetic background impacted IF treatment effects across multiple physiologic systems. In total, over 66,000 body weight measurements, 2,000 frailty assessments, and 1,000 each of hematology, immunology, metabolic phenotype, and glucose tolerance assays were generated by the multi-year study. We performed longitudinal analyses to identify traits significantly influenced by IF treatment as well as to determine which traits exhibited heterogeneity in response to IF by genetic background (**Supplementary Fig. S2**; **Supplementary Tables S6-S10**).

#### Body weight trajectories reveal strain-specific IF responses

Weekly body weight measurements largely demonstrated canonical aging-associated trajectories, with progressive increase to peak mass at maturity followed by gradual decline post-maturity, and either rapid increase or decrease near end of life (n measurements *>*66k; **Figs. 2a and 2b**). Summary metrics—mean mass (MM) and positive area under the curve (+AUC)—showed significant strain-by-diet interaction (p=9.96e-4 and 2.05e-4, respectively, **Supplementary Table S9**), indicating heterogeneous IF responses across genetic backgrounds (**Fig. 2c**). Strains with higher baseline body mass under ad libitum (AL) feeding exhibited greater weight loss under IF (MM: *ρ*=–0.78; +AUC: *ρ*=–0.58). While summary metrics did not exhibit clear sexually dimorphic IF response at the population level (MM sex-by-diet interaction p=0.61, +AUC sex-by-diet interaction p=0.39), the timing of IF effects varied by sex and strain (**Figs. 2a and 2b**). For example, 004/TauUncJ male body weight responded in mid-life but not late-life, and 004/TauUncJ females sustained body weight differences through late life.

**Figure 2:**
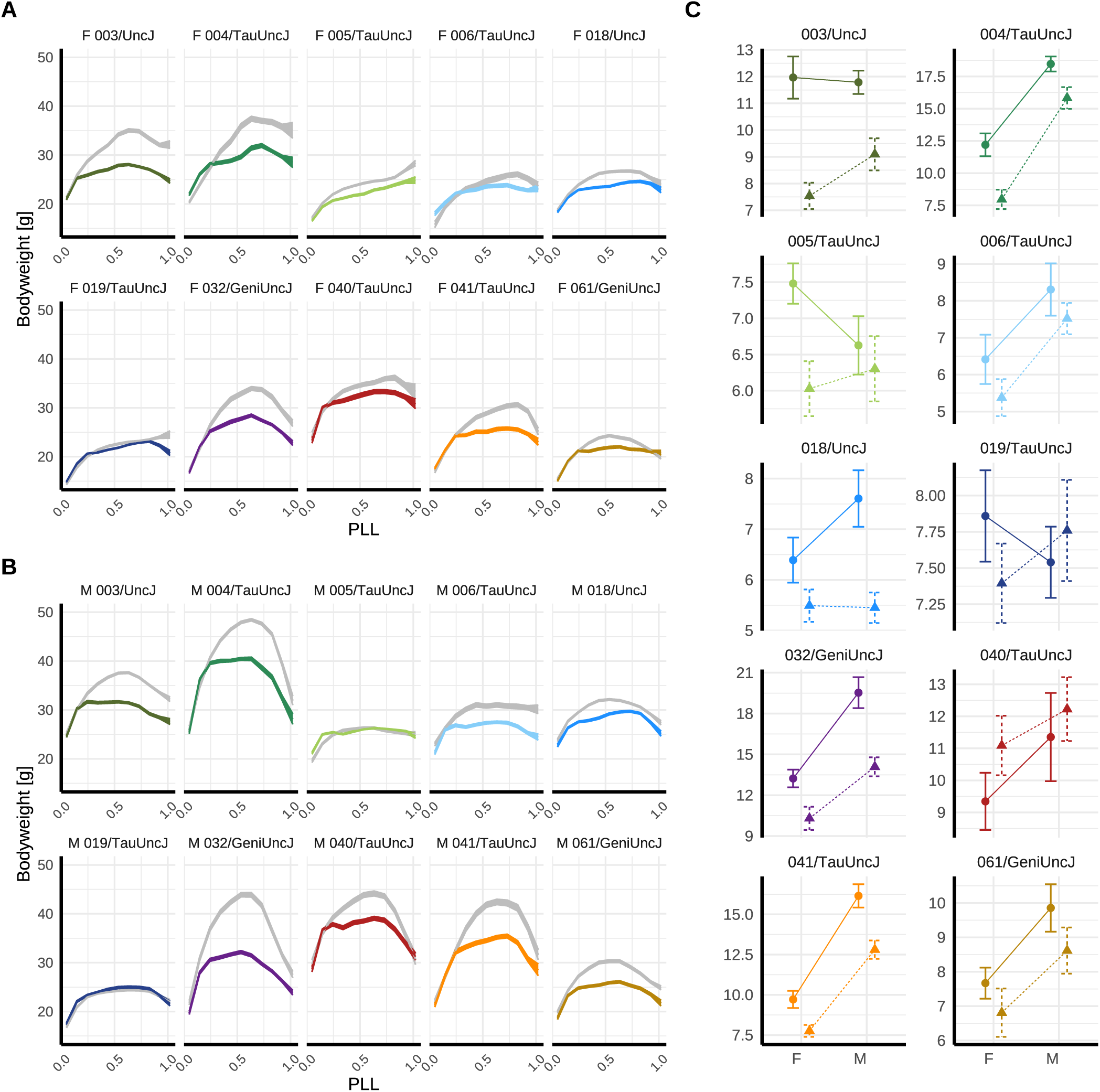
Body weight phenotype response to intermittent fasting (IF) and ad libitum feeding (AL) across 10 inbred strains reveals sexually dimorphic genetic effects. A and B) Longitudinal body weight measurements (n*>*66k) were collected weekly over the lifespan of IF (color) and AL (gray) mice. Mean±SEM body weight (g) was estimated across 20 equidistant spans of scaled age (proportion of life lived, PLL) for (A) females [F] and (B) males [M]. C) Longitudinal body weight measurements collected throughout lifespan were summarized per mouse as total area under the curve for body weight (+AUC[BW]; see Materials and Methods) and plotted as mean±SE +AUC[BW] for male and female mice, grouped by dietary regimen (IF [Δ] or AL [◦]) and genetic strain.

#### Metabolic phenotype effects are ageand strain-dependent

Nuclear magnetic resonance (NMR) body composition analysis provided non-invasive measurements of metabolic profiling, distinguishing lean tissue (organs, muscle) and fat mass components at 10 and 22 months of age, roughly corresponding to middle-aged and older adult mouse life phases (n measurements*>*1,000). NMR-based assessments revealed that IF reduced lean mass in both sexes at 10 months (diet p=8.13e-8 in females; p=4.01e-8 in males) and 22 months (diet p=2.32e-6 in females; p=6.01e-6 in males) but had no significant effect on adiposity at either timepoint (**Supplementary Table S8**). Strain-specific responses were evident for lean mass (strain-by-diet variance p=1.21e-4) and adiposity (p=2.36e-7) (**Fig. 3**, **Supplementary Fig. S3**). Strains with greater lean mass and adiposity loss under IF tended to exhibit higher respective baseline values under AL feeding (lean mass: *ρ*=–0.58; adiposity: *ρ*=–0.57). However, 019/TauUncJ mice lost lean mass while gaining adiposity across both ages and sexes, underscoring the importance of composition-specific metrics for characterizing metabolic phenotype.

**Figure 3:**
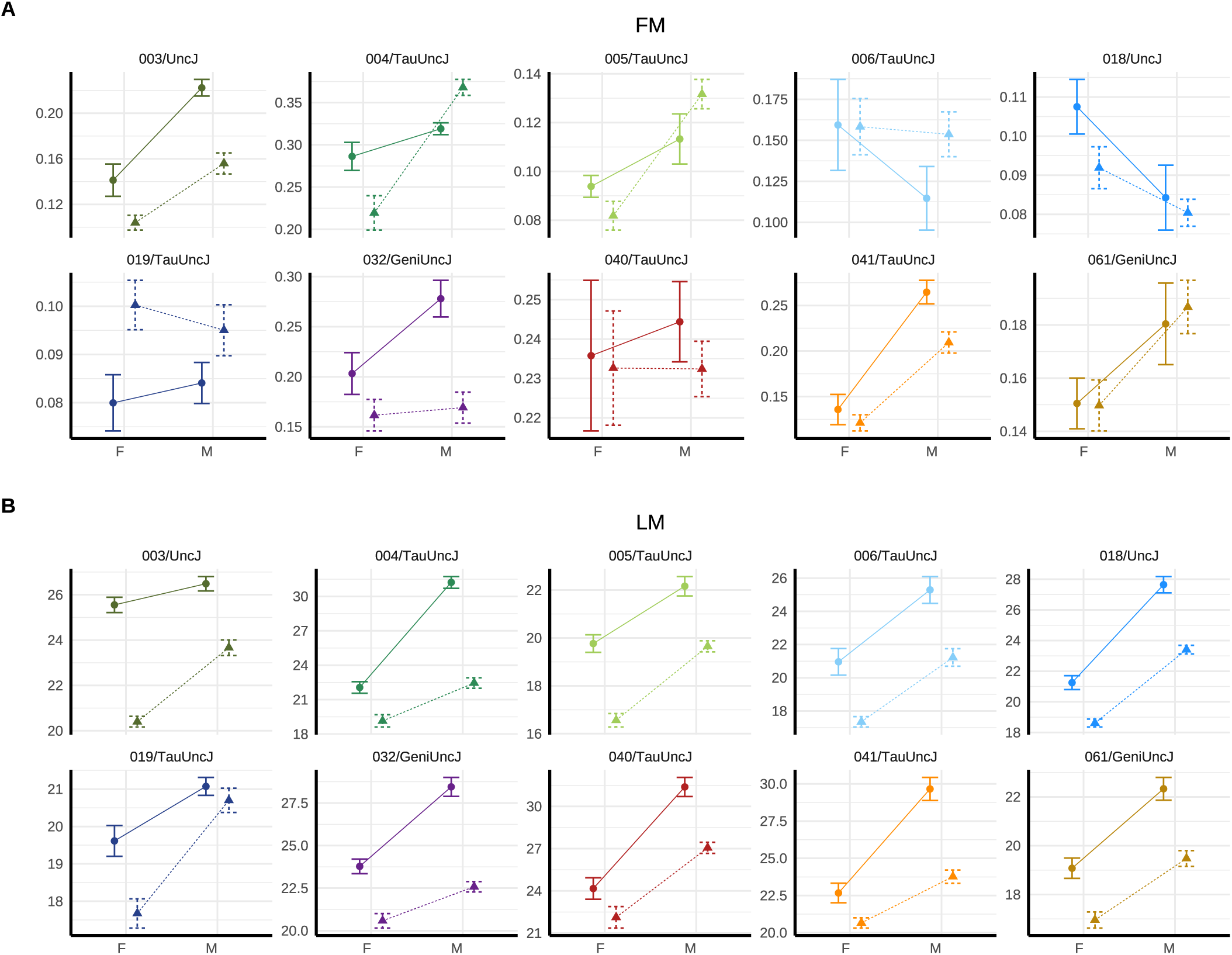
Metabolic phenotype response to intermittent fasting (IF) is influenced by genetic background. Nuclear magnetic resonance (NMR) body composition assays were conducted at 10 and 22 months (n*>*1k). Month 10 NMR data were plotted as (A) mean±SE adiposity [%], a measure of fat mass, and (B) lean mass [LM, g] for male and female mice–grouped by dietary regimen (IF [Δ] or AL [◦]) and genetic strain. Early lean mass loss in response to IF was observed with varied risk across strains and sexes. Some strains (e.g., 005/TauUncJ, 006/TauUncJ) showed minimal total mass change but significant lean mass loss, underscoring the importance of composition-specific metrics for characterizing metabolic phenotype. Plots for 22 months shown in **Supplementary Fig. S3a-b.**

#### Frailty trajectories vary by genotype but not diet

Multisystem frailty was determined by a set of 27 serially collected non-invasive biomarkers of frailty (weeks 21 [preintervention], 43 [intervention onset], 95, 121, 147; total n frailty index [FI] assessments*>*2,000). Per-mouse frailty trajectories were summarized as linear coefficients (slopes) on proportion of life lived (PLL [scaled age]) to describe pace of aging, and final FI score was used as a proxy for lifetime multisystem frailty accumulation. Neither the rate of frailty accumulation nor final frailty scores differed significantly between IF and AL groups (**Supplementary Table S8**). Further, statistical models did not support strain-specific dietary response. Simplified models without strain-specific dietary response showed strong strain effects (strain variance p*<*2e-16 for final frailty score and p=9.25e-3 for rate of frailty accumulation). Together, these results indicate genetic background modulates vulnerability to frailty, and strain-specific normative aging as measured by frailty index is unperturbed by IF in the CC panel studied.

#### Lifetime incidence of select health deficits altered by IF in strain-dependent manner

Although frailty is often reported as an aggregate expression of risk resulting from accumulation of health deficits across multiple physiologic systems [23–25], we also explored more fine-grained evidence of strain-specific aging by comparing cumulative incidence of individual health deficits (**Fig. 4**, **Supplementary Fig. S4**). Regardless of experimental condition or genetic strain, most indicators of aging were infrequently observed: breathing rate/depth, cataracts, corneal opacity, dermatitis, diarrhea, eye discharge or swelling, gait disorders, malocclusions, nasal discharge, rectal prolapse, righting reflex, tail stiffening, tremor, tumors, and genital prolapse. Alopecia and microphthalmia were also infrequently observed but there were exceptional strains. For example, CC018/UncJ was both more susceptible to alopecia in the control condition and showed stronger IF response than other strains.

**Figure 4:**
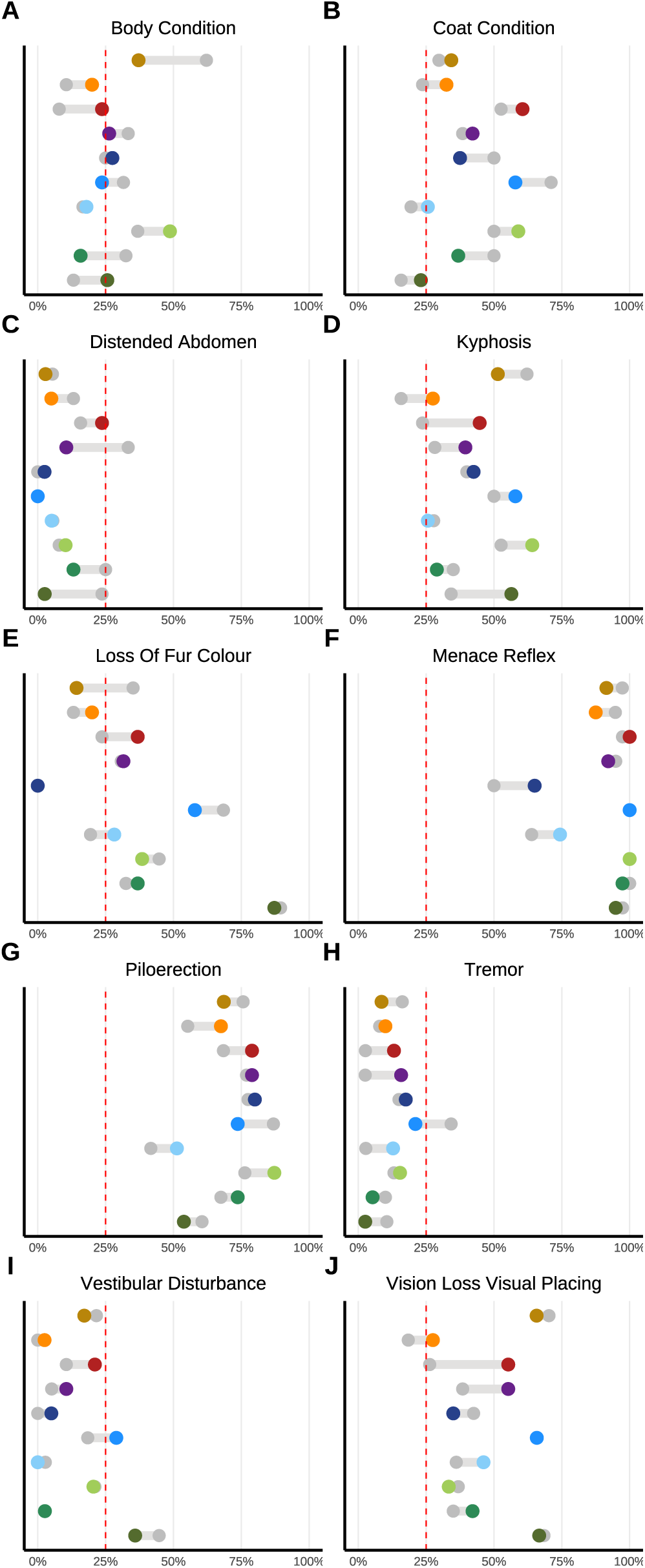
Genetic and dietary regulation of lifetime-accumulated health deficits across multiple physiological systems. Twenty-seven serially collected non-invasive biomarkers of frailty were measured at weeks 21 (preintervention), 43 (intervention onset), 95, 121, and 147 weeks (n*>*2k). The comparison of cumulative incidence of individual health deficits across intervention groups (IF = color, AL = gray) reveals distinct aging patterns specific to each strain. Note: Remaining items shown in **Supplementary Fig. S4**. Frailty index items were binarized as severe/nonsevere.

Subtracting per-strain incidence among the remaining 10 indicators demonstrated varied IF response across genetic backgrounds. For instance, CC003/UncJ was particularly susceptible to loss of fur color yet showed little response to IF, and IF protocol assignment was associated with a 20% higher risk for displaying menace reflex in CC0019/TauUncJ. CC018/UncJ was vulnerable to diet effects for several of the more commonly observed items. For strains with meaningful evidence of IF effect on kyphosis phenotype, kyphosis was more commonly observed in mice on IF treatment than controls, although extent of this adverse effect varied, and cumulative incidence remained below 50% for all except CC005/TauUncJ mice on treatment.

#### Hematologic aging markers reveal complex effects of IF

We conducted hemato-phenotyping longitudinally to characterize whole blood cell type composition at 10 and 22 months (n assessments*>*1,000). Complete blood counts showed significantly elevated red cell distribution width (RDW-CV) in IF-treated mice in both sexes at mid-(p=1.64e-5 in females, p=8.88e-5 in males) and late-life (p=2.1e-4 in females, p=1.63e-4 in males), consistent with erythropoietic stress. In contrast, mean corpuscular volume (MCV) did not differ significantly between IF and AL groups in either sex at either time point (**Supplementary Table S8**), indicating that average red blood cell size remained within normative strain- and sex-specific ranges. The combination of elevated RDW and stable MCV is consistent with early-stage or mixed anemia. This profile may reflect mild nutritional deficiencies during fasting (e.g., iron or B12 deficiency) or mixed anemia, in which microcytic and macrocytic populations offset to yield a normal MCV. Strain-specific responses were evident for both RDW (strain-by-diet variance p=2.67e-18) (**Fig. 5a**, **Supplementary Fig. S5a**) and MCV (p=0.0018) (**Fig. 5b**, **Supplementary Fig. S5b**). Strains with greater RDW and MCV elevation under IF tended to exhibit lower respective baseline values under AL feeding (RDW: *ρ*=–0.66; MCV: *ρ*=–0.76).

**Figure 5:**
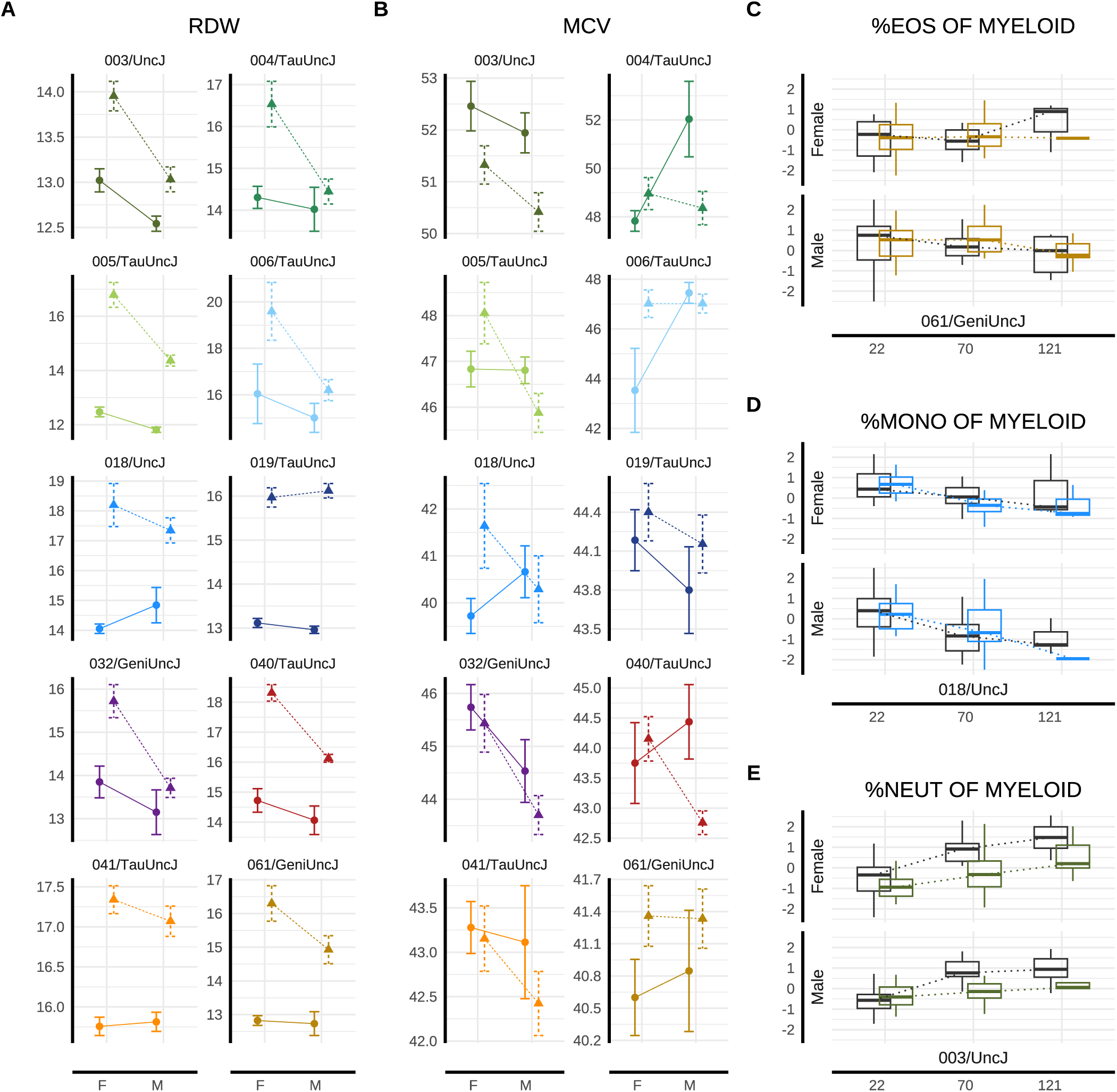
Genetic signatures of hematologic and immunologic response to IF. Longitudinally-collected data for representative hematologic and immunologic health outcomes are presented. Year 1 red blood cell distribution width (RDW CV) data (A) and mean corpuscular volume (MCV) (B) were plotted as mean±SE for male and female mice grouped by dietary regimen (IF [Δ] or AL [◦]) and genetic strain, demonstrating sexually dimorphic IF response at 10 months. Longitudinal immunologic outcomes are presented across randomly sampled genetic backgrounds and both sexes at mid- and late-life for eosinophil (C), monocyte (D), and neutrophil/granulocyte (E) concentration among myeloid cells (IF = color, AL = gray). Strain colors are consistent across all figures and follow the key provided in 5a. Of the three myeloid cell subtypes, only monocyte abundance showed statistical evidence of strain-by-diet interaction in this study. Note: 22-month data for RDW and MCV are shown in **Supplementary** Fig. 5.

From the broader hematologic panel, hemoglobin concentration, red blood cell (RBC) count, and reticulocyte levels are particularly informative to further characterizing the anemia-like phenotype. At 10 months, IF was associated with sex-specific alterations: females exhibited reduced hemoglobin (p=0.015) and RBC counts (p=0.006); no significant differences were observed in males or older females. Reticulocyte counts were elevated in both sexes at 10 months (p=4.42e-4 in females, p=0.004 in males) but not 22 months (**Supplementary Table S8**), suggesting early compensatory erythropoietic response.

### Mouse models of hematologic response

To further investigate the structure of hematological trait variation in response to IF across sex and time, we visualized strain-, sex-, and timepoint-specific diet effect scores (see Methods: Mouse model selection). The resulting heatmaps (**Supplementary Fig. S6**) revealed patterns in IF hematologic response profiles. Hematologic traits were partitioned into clusters based on IF responses in year 1 females, and row order was preserved across all panels to facilitate cross-stratum comparisons. Three major clusters emerged for hematologic trait response to IF: (1) erythroid and lymphoid morphology and abundance; (2) granulocyte and monocyte populations alongside red cell concentration metrics; and (3) red cell distribution and reticulocyte indices, reflecting coordinated and biologically plausible shifts in hematopoietic activity in response to IF.

Strain clusters of hematologic response profiles varied by sex and timepoint, with some strains showing consistent profiles across study strata and others displaying sex- or timepoint-specific divergence. For instance, by year 1 (midlife), four strains (003/UncJ, 004/TauUncJ, 041/TauUncJ, and 006/TauUncJ) showed similar hematologic profiles within males and within females—e.g., lesser vulnerability to high RDW on IF. Within males, strains 032/GeniUncJ and 040/TauUncJ showed similar patterns to these four strains’ midlife hematologic IF response but were distinctive from the strain/model cluster in females, suggesting mouse model development for IF response will be sex-specific. Clustering patterns were mirrored between year 1 males and year 2 females, and vice versa, further indicating complex, time-dependent strain effects on hematologic phenotypes.

#### IF induces sex-specific remodeling of immune cell composition across strains

We conducted immuno-phenotyping longitudinally to characterize and classify various immune cell subtypes (T cells, B cells, natural killer (NK) cells, dendritic cells, and myeloid cells) and their proportions longitudinally via flow cytometry (n assessments*>*1,000). Flow cytometry profiled major immune cell subsets pre-intervention at 5 (pre-intervention) and 16 and 28 months (post-intervention). Immune cell subsets included T cells, B cells, natural killer (NK) cells, dendritic cells, and myeloid cells. By 16 months, IF elicited distinct sex-specific differences in IF response across both lymphoid and myeloid compartments, as detailed below. Low survival to the 28-month endpoint precluded meaningful analysis.

Within the lymphoid lineage, B cell frequencies were significantly altered under IF in males (p=0.026) but not in females (p=0.972) at 10 months following intervention onset. CD4^+^ T cells also exhibited male-specific differences in IF response (p=0.001), with no significant change in females (**Supplementary Table S8**). NK cells appeared particularly responsive to long-term dietary modulation, potentially reflecting innate immune adaptation to metabolic stress. NK cells (as a proportion of lymphocytes) were significantly altered by IF in both sexes, with a more pronounced effect in females (p=7.62e-6) than in males (p=1.44e-3). Strain-specific responses were not evident for these lymphoid lineage cells (**Supplementary Table S9**), suggesting low genetic modulation of IF responsiveness among lymphoid immune traits.

Within the myeloid lineage, eosinophil proportions were significantly altered in response to IF among both females (p=0.018) and males (p=1.37e-4). Neutrophils, key innate responders to infection or inflammation, also showed significant response to IF in both sexes (females: p=1.74e-3; males: p=6.03e-5). In contrast, monocyte abundance (among myeloid cells) was significantly altered under IF only in males (p=2.27e-3; **Supplementary Table S8**). Of these myeloid subsets, monocyte abundance was the only trait to exhibit a significant strain-by-diet interaction (p=1.99e-3; **Supplementary Table S9**), suggesting low genetic modulation of IF responsiveness among myeloid immune traits (**Figs. 5c to 5e**).

### Mouse models of immunologic response

To further characterize the structure of immunologic response to IF, we visualized strain-, sex-, and timepoint-specific diet effect scores (see Methods: Mouse model selection). Heatmaps of IF scores (**Supplementary Fig. S7**) revealed patterns in strain-specific IF immune response profiles. Immunologic traits were clustered based on IF responses in year 1 females, and row order was preserved across all panels to facilitate cross-stratum comparisons. Three major clusters emerged: (1) myeloid and lymphoid cell abundance and activation (2) a lymphocyte-dominant cluster; and (3) a cluster dominated by inflammatory and innate immune markers, reflecting coordinated and biologically plausible shifts in immunologic activity in response to IF. Three strains (003/UncJ, 061/GeniUncJ, and 019/TauUncJ) may possess shared regulatory architectures governing IF-induced immune modulation, as their immune response profiles cluster together within all sexes and timepoints.

### Comparative analysis finds phenotypic response to IF differs between inbred and outbred mice

We previously reported longitudinal phenotype data from the DRiDO study for female Diversity Outbred (DO) mice including AL and 2-day IF cohorts [9]. We performed longitudinal analysis in a subset of these data (mice assigned to IF or AL) to determine which traits exhibited heterogeneity in response to IF across highly diverse outbred population (**Supplementary Table S11**). Despite nearly identical protocols and husbandry conditions, IF elicited both shared and divergent physiological responses in outbred (DO) and inbred (CC) mouse populations.

#### Body composition trajectories diverge by genetic background

IF significantly reduced lifetime body mass in both populations (outbred +AUC p=3.96e-14; inbred +AUC p=3.2e-3), with comparable effect sizes. However, lean mass responses diverged: outbred females maintained lean mass under IF (**Supplementary Table S11**), while inbred females showed significant reductions (diet p=8.12e-8 at 10 months; p=2.32e-6 at 23 months, **Supplementary Table S8**), suggesting strain-specific susceptibility to sarcopenia. Population-level resilience to lean mass loss among outbred females with high genetic diversity may have obscured the potential for genetic predisposition to loss of lean mass on IF observed in inbred females.

#### Frailty outcomes reflect genetic context, extended lifespan in outbred mice

In outbred mice, IF reduced cumulative frailty at end of life (p=0.0024) without significantly altering the rate of frailty accumulation (**Supplementary Table S11**), indicating improved late-life healthspan without a change in biological aging rate. In the inbred panel, frailty outcomes varied by strain but were similar across diet groups. Longer lifespan in outbred mice may have extended the window for frailty accumulation, complicating direct comparisons.

#### Hematological responses to IF are highly concordant across recombinant inbred and outbred cohorts

IF increased red cell distribution width (RDW-CV), a biomarker of erythropoietic stress and aging, across both outbred and inbred populations (outbred: p*<*2e-16 at 10 months and 22 months; inbred: p=1.64e-5 at 10 months, p=2.10e-4 at 22 months). This robust, genotype-independent effect supports RDW-CV as a candidate biomarker of adverse IF response.

#### Immune cell profiles show population-specific effects

By 16 months, IF significantly increased lymphocyte proportions (of viable cells) in outbred mice (p=3.15e-7, **Supplementary Table S11**), whereas evidence was inconclusive in inbred females (**Supplementary Table S8**). IF also reduced the proportion of effector CD4^+^ T cells (as fraction of total CD4^+^ T cells) in outbred mice at this timepoint (p=4.89e-4, **Supplementary Table S11**); effects were not statistically significant among female inbred mice (**Supplementary Table S8**). The absence of significant immunologic findings in inbred female CC mice among traits selected for strong IF effects in outbred female DOs may reflect reduced genetic diversity in the recombinant panel.

### Thermoregulatory decline in late life

Generalized additive modeling was used to examine how body temperature varies with sex, strain, body weight at six months, and scaled age, while accounting for repeated measures through a random effect for mouse identity [see Methods: Non-linear effects]. Scaled age and body weight were modeled using smooth terms to capture nonlinear effects. Temperature exhibited a well-characterized nonlinear trajectory with age, with an inflection point occurring around 80% of the lifespan, consistent with prior observations of thermoregulatory decline in late life (**Fig. 6**). To assess the contribution of diet, a nested model comparison was performed using an analysis of deviance. The addition of diet did not significantly improve model fit (p=0.386), indicating that dietary effects are negligible in this context. Visually apparent variation in temperature patterned by sex and genetic background was not significantly associated with these factors, contrary to expectations regarding the dominant role of these intrinsic biological factors in shaping thermoregulatory dynamics.

**Figure 6:**
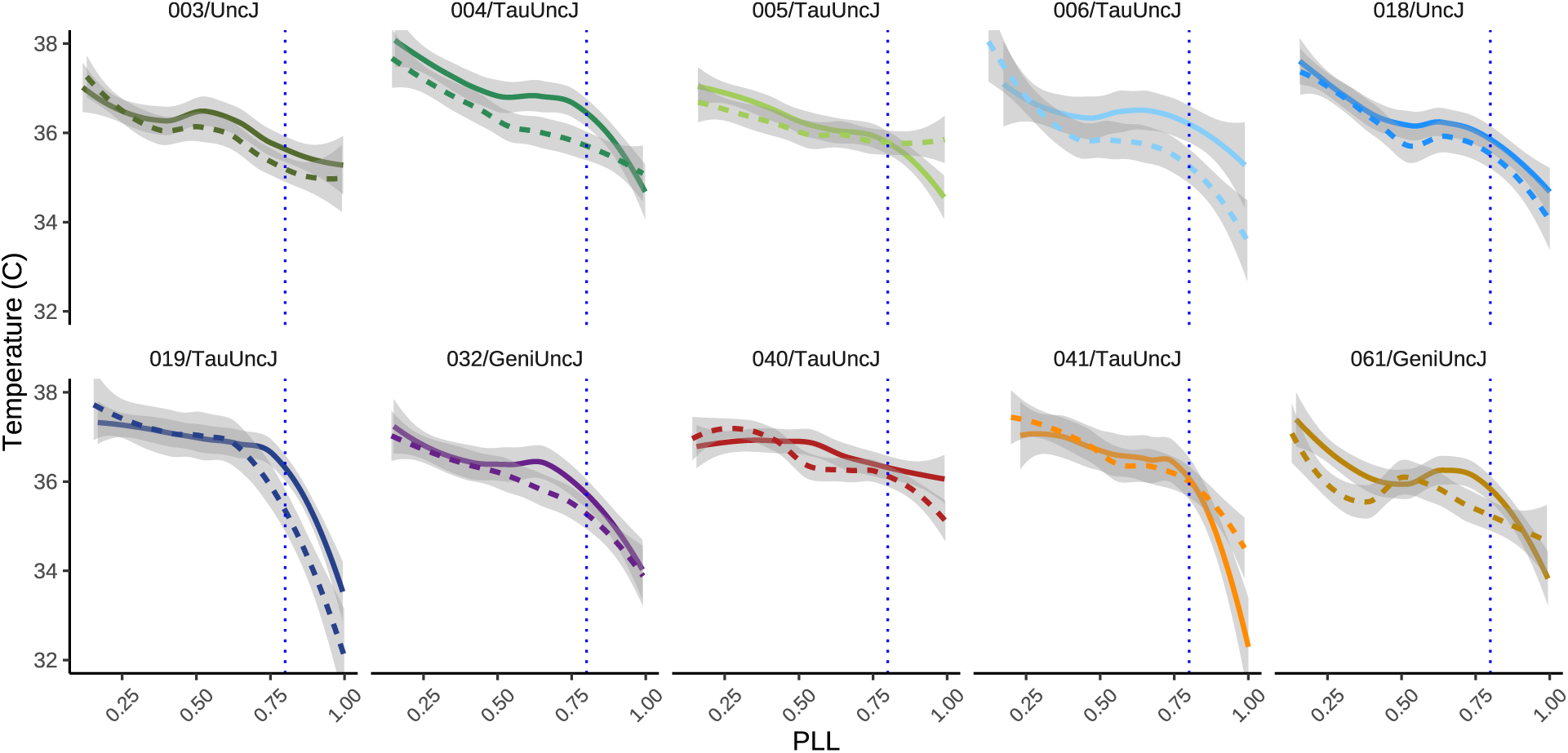
Inflection point in body temperature at late-life transition. Body temperature plotted against proportion of life lived (PLL) revealed a distinct inflection point at approximately 80% PLL for most genetic strains. This late-life shift suggests a physiological transition that may mark the onset of age-related decline or altered thermoregulatory control. Data are shown across the lifespan, with temperature values averaged within PLL bins. Note: solid line=female, dotted line=male.

### Longevity biomarkers

Analysis of biomarkers underlying heterogeneous longevity within intervention groups reveals several key findings (**Fig. 7**, **Supplementary Table S12**). First, greater IF-induced phenotypic changes in health and metabolic traits do not necessarily lead to lifespan extension relative to sex-, strain-, and genotype-matched peers. This observation replicates patterns from our parallel study of outbred females (Di Francesco et al. 2024) and extends these findings to inbred mice of both sexes. Second, late life adiposity was associated with higher lifespan after adjustment, indicating a protective longevity effect for mice that retained relatively higher proportions of adipose tissue within sex, strain, and diet group. Third, intra-individual homogeneity in red blood cells (here, hemoglobin concentration) is strongly associated with lifespan extension, more so than other hematologic traits. Fourth, both innate and adaptive immune cell types play significant roles in inter-individual lifespan variation. Longer-lived mice exhibited lower levels of CD11b^+^ myeloid cells in mid-life, suggesting reduced innate immune activation during this period may support longevity. In contrast, higher levels of eosinophils (as a proportion of myeloid cells) in mid-life, and higher lymphocyte and B cell abundance (as proportion of viable cells and lymphocytes, respectively) in late-life, were associated with longer lifespan. These findings indicate that increased lymphocyte abundance and altered myeloid cell proportions are correlated with healthier aging at distinct life stages. Although causality remains to be determined—potentially through cell depletion studies or immune challenge models—the observed immune shifts may reflect mechanisms such as reduced inflammaging and enhanced immune memory or cancer surveillance.

**Figure 7:**
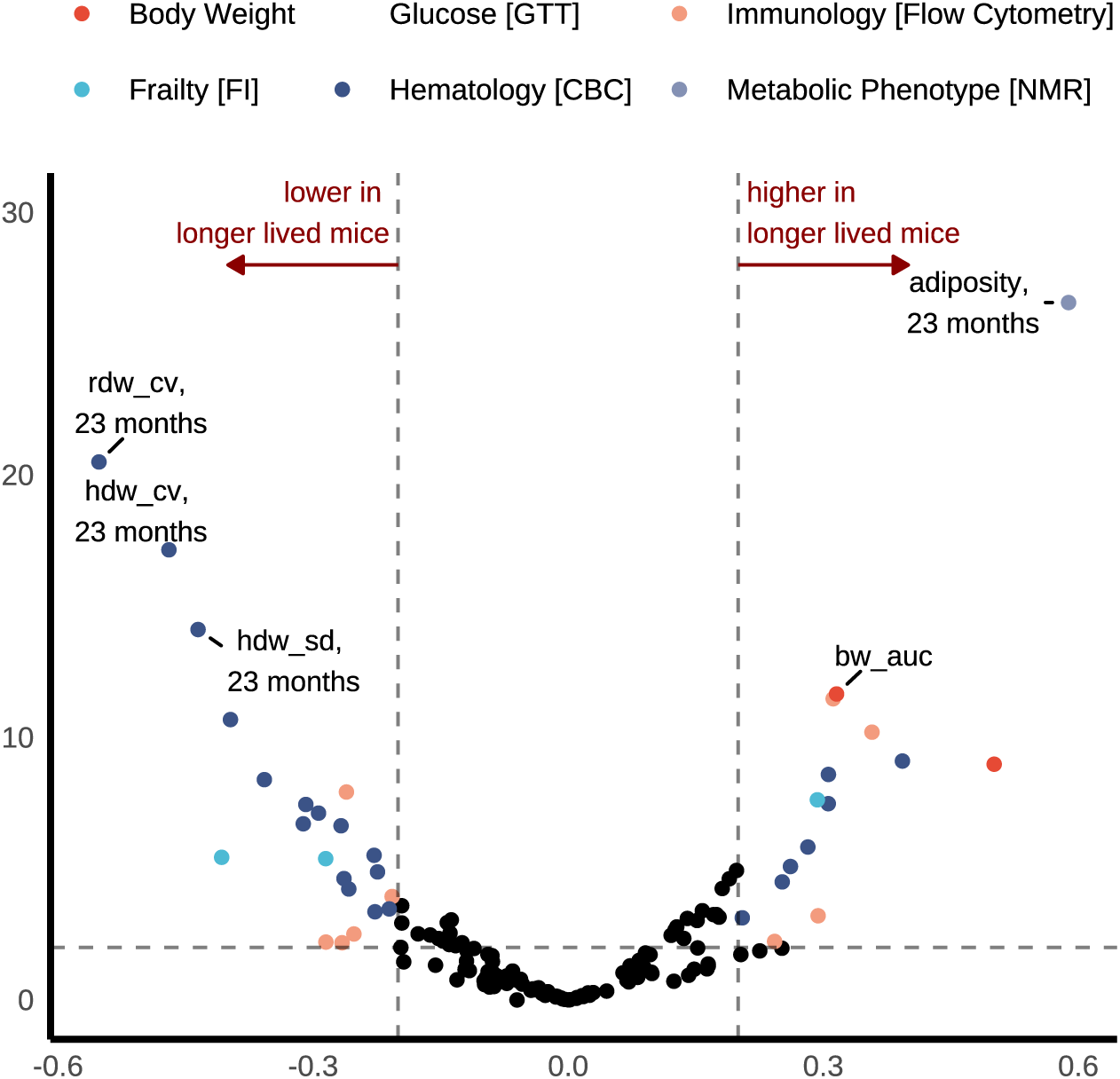
Comprehensive longitudinal phenotyping identifies biomarkers for longevity underlying heterogeneous IF effects on lifespan within strain and sex. Regression analysis on lifespan with traits at each timepoint after adjusting for effects of diet, sex, strain, batch, and 6-month body weight. All continuous variables were rank normalized before model fitting, resulting in standardized beta coefficients (x-axis). We performed likelihood ratio tests to obtain p-values for the adjusted associations. FDR adjustment accounted for multiple testing. Traits are categorized and presented in volcano plots by physiologic domain. Results largely cohered with longevity biomarkers identified among female Diversity Outbred mice in our parallel study DRiDO. Weekly body weight outcome was modeled as MM and +AUC (defined above).

## Discussion

In the current study, we leveraged the genetically diverse and reproducible Collaborative Cross recombinant inbred strains to characterize genetic heterogeneity in response to 2-day intermittent fasting treatment. We found modest support for the presence of genotype-by-treatment (GxT) effects on lifespan as well more substantial GxT effects on body weight, adiposity, and hematological traits including red cell distribution width (RDW). These outcomes support the hypothesis that response to dietary interventions, specifically IF, is genetically dependent and variable in an animal model with similar physiological features to humans.

### Sexual dimorphism in longevity intervention response

Sexual dimorphism of genotype effects on phenotypic traits has been observed in large-scale mouse studies such as the International Mouse Phenotyping Consortium (IMPC) database (*>*14k wildtype animals and *>*40k mutant mice) [26] and the Intervention Testing Program [27, 28]. The latter also revealed unexpected sex differences in geroprotective effects of some interventions (e.g., [29, 30]). In keeping with our findings, dietary treatments have been shown to affect lifespan differently in males and females [11, 14, 31]. These findings may reflect sex-specific dietary requirements for healthy aging that could not be explored in our parallel study of female outbred mice.

### Lifespan response to IF differs between inbred and outbred mice

We previously reported survival data from the DRiDO study for female Diversity Outbred (DO) mice including AL and 2-day IF cohorts [9]. The DO are derived from the same founders as the CC strains and thus share the same genetic variants. In addition, the two studies were carried out concurrently in the same mouse facilities. In the DRiDO study, which included only female mice, the 2-day IF intervention led to significant lifespan extension – population-level median lifespan increased by months. We further observed that lifespan increase was more pronounced in mice with lower pre-intervention body weight and no extension of lifespan was observed for the heaviest mice. In the CC study, aggregate lifespan data revealed sexually dimorphic lifespan response to IF. We observed significant lifespan extension in aggregated (across-strain) data for male mice only, which is discordant from the result in outbred mice.

### Heritability of lifespan is concordant in inbred CC and outbred DO mice

It is common in preclinical studies to test intervention effects on a single inbred mouse strain, typically C57BL/6. Yet lifespan is a complex trait that is influenced by a large number of genetic and non-genetic factors that drive phenotype heterogeneity. Estimates of heritability provide some information on the relative importance of genetic factors in comparison with non-shared environmental factors. In both the CC Longitudinal Study and DRiDO, we imposed precise control over the environment and husbandry of mice to minimize environmental variance and selected mouse models with exceptionally high levels of diversity to maximize the genetic contribution thereby creating the most favorable setting for high heritability of lifespan and other phenotypes. As in DRiDO, we found genetic background proved to be the more important factor in determining lifespan. Heritability of lifespan in the CC study was nearly completely accounted for by additive genetic variance—ie., variance due to mean effects of single alleles. We estimated the narrow and broad heritabilities for lifespan (adjusted for sex and diet) as being nearly equal. This does not preclude the possibility of gene interactions affecting lifespan and IF response as traits with large additive variance can be driven by physiological epistasis [32].

### Genetics alter phenotypic response to intermittent fasting

We identified gene-by-treatment interaction across metabolic, hematologic, and immunologic profiles. Shifts from strain-normative phenotype were often distinguishable by both genetic strain and sex. Intervention studies that incoporate genetic variability will be important for success in translational metabolic research. The Collaborative Cross (CC) study presented here demonstrates that dietary intervention response is nonuniform and patterned across genetic backgrounds. By profiling intermittent fasting effects across ten inbred strains and multiple physiological systems, this study establishes a reusable library of mouse models for mechanistic follow-up, enabling validation of genetically driven heterogeneity in intervention outcomes previously observed among outbred DO mice. Future studies using these mouse models could regenerate the same genotypes to replicate our findings as well as exploring new genetic backgrounds, including new CC strains and their F1 hybrid to further explore the variability in dietary intervention outcomes.

### Concordance and divergence in the impacts of intermittent fasting on health among genetically diverse inbred and outbred mice

The Collaborative Cross (CC) Longitudinal Study, which explores gene-by-treatment interaction in a panel of recombinant inbred strains, was conducted in parallel with DRiDO, a study of dietary restriction (including 2-day IF) in female DO mice. Uniquely, the two studies were conducted in the same laboratory facility during the same time period. IF produced broadly concordant effects on health across genetically diverse mouse populations, with some notable divergences. In both the outbred (DRiDO) and inbred (CC) studies, IF significantly reduced lifetime body mass in females, though strain-specific deviations in late-life weight trajectories were observed among certain inbred lines. Lean mass was preserved in outbred females on IF, contrasting with reductions seen across the recombinant inbred strains. IF elevated red cell distribution width (RDW-CV) across both cohorts, highlighting a robust, genotype-independent biomarker response warranting mechanistic investigation, especially in light of consistently strong predictive utility as a longevity biomarker (see Results: Longevity Biomarkers). Leveraging both outbred and recombinant inbred mouse populations provides complementary strengths for dissecting the biology of dietary interventions. Outbred populations, like those used in the DRiDO study, are powerful for genetic mapping and broadly representative of genetic diversity, a strategy increasingly adopted across model organisms since its formalization in the early 2000s [33]. In contrast, the recombinant inbred panel screen used here—less commonly employed—offers a replicable framework for identifying strain-specific responses.

### Inflection point in thermoregulatory response

A rapid decline in body temperature (TB) is an indicator of immanent death in standard laboratory mice (e.g., C57BL/6) [34]. The predictive power of TB was recently confirmed in DO mice [35]. In the present study we sampled TB at 5 timepoints and rescaled chronological age to proportion of life lived. We confirmed the expected decline in TB with age, and observed an inflection point at 80% of life lived (80 PLL) that was most pronounced in strains CC004, CC006, CC019, and CC041. The non-linear rate of change as a function of PLL was not impacted by IF. These trends are apparent at the strain-average level, but at the individual level can be obscured by noise (measurement error) or limited time density of sampling (6-month intervals). These finding support TB as a biomarker of aging and predictor of lifespan that is consistent across genetic background.

### Comprehensive longitudinal phenotyping identifies longevity biomarkers

In our previous work [9], health and metabolic traits were sensitive to diet but were poor predictors of lifespan, while hematologic and immune traits showed strong dietary response and lifespan prediction. That is, counterintuitively, variability in lifespan within treatment group was not explainable by favorable changes in response to dietary restriction. We looked for replications of these patterns in the CC, considering each phenotypic domain and its relevance as a marker for longevity. As in DRiDO, the current study found that IF-induced phenotype changes in health and metabolic traits did not necessarily translate into lifespan extension. In light of these contradictory effects, and the adoption of fasting by people who seek to improve health and increase lifespan, further study of health effects of IF should be a priority for future research.

A recent trial [36] found that despite improved human adherence to IF resulting in significant body weight and fat loss, clinically meaningful improvements in systemic inflammatory markers or glucose-insulin metabolism were not observed. These findings further validate that weight reduction does not always translate into improved metabolic heath and highlights a priority area for future research on IF as an anti-aging intervention.

### Computational methods for recombinant inbred strain panel screens

Recombinant inbred panel screens, such as the one presented here, require large sample sizes to achieve sufficient statistical power for detecting disaggregated strain-specific effects. This scale of experimentation is often impractical. Computational methods can be leveraged to maximize power to detect variability in strain-specific intervention response. In this study, we introduced a novel application of empirical Bayes estimation of strain-specific random effects to identify mouse models exhibiting heterogeneous responses to intervention. This approach conceptualizes strain in the panel screen as samples drawn from a broader population, enabling genotype-by-treatment (GxT) interactions to be modeled as a single variance component representing intervention effect variability across genetic backgrounds. The magnitude of this variance component can be compared across models of standardized phenotypes to identify traits more pronounced GxT interaction effects. Further, by leveraging empirical Bayes estimates, distinctive strains can be identified by examining the extreme values of the best linear unbiased predictors (BLUPs) for strain-specific intervention effects. E.g., selecting strains with maximum and minimum BLUPs for dietary effect enables the selection of strain pairs that exhibit divergent, and potentially directionally opposite, responses to the intervention.

### Limitations

This study aimed to investigate heterogeneity of lifespan extension in response to IF across a panel of 10 recombinant inbred strains. However, the observed effect of IF on lifespan was modest relative to the within-strain variance, limiting our ability to establish strong conclusions about strain-specific differences in response. Future studies may benefit from examining genotype-by-treatment (GxT) interactions using dietary interventions with more pronounced effects on lifespan—such as 40% caloric restriction—though IF remains of high interest due to its tolerability [37].

Another limitation lies in the study’s sample composition. The 10 strains included were selected based on availability within the study’s timeframe, constituting a convenience sample. While strain was modeled as a random effect, the representativeness of these strains relative to the broader Collaborative Cross (CC) population is uncertain. Ideally, future studies would incorporate a larger and more diverse set of strains to better capture the genetic landscape of response variability. However, scaling up the number of strains and/or the number of animals per strain poses significant logistical and resource challenges, underscoring the need for more efficient study designs in lifespan research. To fully harness the potential of multiparental recombinant inbred panels such as the CC in advancing scientific discovery, continued innovation in experimental design is needed.

Future studies with more densely sampled time series data will avoid data sparsity issues across the full range of PLL. We were able to demonstrate this method for temperature, which was more frequently collected than other phenotypes (5 measurements vs. 2 to 3). Normalizing chronological age across all phenotypes could reveal non-linear dynamics in IF response. Such studies are in the pipeline.

Despite these limitations, the study design proved effective in detecting robust treatment responses in other phenotypes. We observed significant and substantial heterogeneity in body weight and composition responses to IF, suggesting that the design is well-suited for outcomes with stronger intervention effects. These findings highlight the importance of aligning study design with the expected magnitude of treatment effects when investigating complex traits across genetically diverse populations.

## Conclusion

This study has demonstrated that the diverse genetic backgrounds available in the Collaborative Cross inbred strain panel induce variable IF treatment response across multiple health- and aging-related phenotypes. These outcomes support the finding that dietary treatment response is genetically dependent and variable in an animal model with similar physiological features to humans. Genetic diversity in the study addresses common limitations of single-genome research, allowing researchers to explore intervention effects across a broad genetic landscape, improving the generalizability and translational potential of findings.

## Materials and Methods

### Animals

Approval for this study was obtained from The Jackson Laboratory Institutional Animal Care and Use Committee. All methods were performed in accordance with the guidelines and regulations of the Committee. The current study included 800 mice (400 females and 400 males), evenly distributed across 10 Collaborative Cross strains (CC003/UncJ, CC004/TauUncJ, CC005/TauUncJ, CC006/TauUncJ, CC018/UncJ, CC019/TauUncJ, CC032/GeniUncJ, CC040/TauUncJ, CC041/TauUncJ, and CC061/GeniUncJ) sourced from the Jackson Laboratory. Mice were housed in a room maintained at 70*^◦^*±2*^◦^*F on a 12/12 h light/dark cycle from 6:00 AM to 6:00 PM and fed a standard chow diet (5K0G, LabDiet). Mice were observed until natural death or ethically mandated euthanasia to acquire full lifespan data. The mice lived their lifespan without handling except for cage changes every other week, scheduled feeding, and phenotyping.

### Diet

All mice were maintained on ad libitum feeding diet until 6 months of age. From six months of age, mice that were randomized to the AL diet had unlimited food access; fresh food was provided weekly when cages were changed. In rare instances when the AL mice consumed all food before the end of the week, the food was topped off mid-week. For mice randomized to the IF diet, 48 hours of fasting was imposed weekly from Wednesday noon to Friday noon. Mice on IF treatment were provided unlimited food access (similar to AL mice) on their non-fasting days. The IF regimen mimicked long periods of food deprivation that are typically experienced by animals in the wild [38]. Mice were maintained on IF or AL per random assignment for the duration of their natural lifespan.

### Phenotyping

All assays were conducted at The Jackson Laboratory following standard operating procedures. Multiple assays were performed and repeated throughout the lifespan to assess physiological status.

#### Body weight

Body weight was assessed weekly, resulting in tens of thousands of longitudinally collected body weight values. Weekly body weights were analyzed after local polynomial regression fitting to one third of data nearest the fitted value within mouse (i.e., loess smoothing).

#### Metabolic Phenotype

NMR body composition analysis was performed at approximately 12-month intervals for each animal, providing noninvasive measurements of fat and lean mass, total body water and free water (grams) via the Echo MRI (Houston, TX) instrument.

#### Frailty and Body Temperature

Frailty data were collected longitudinally at 21 weeks (pre-intervention), at 43 weeks (intervention onset), and concurrent with intervention at 68-, 95-, 120-, and 147-weeks of age (i.e., biannually). Frailty items were scored on a two-[0,1] or three-[0,0.5,1] level ordinal scale where 0 indicated the absence of the health status deficit; 0.5 indicated mild deficit; and 1 indicated severe deficit. Temperature was concurrently collected.

#### Hematologic Traits

We measured hematologic traits in peripheral blood at approximately 12-month intervals for each animal at 45-, 97-, and 149-weeks of age. Blood samples were run on the Siemens ADVIA 2120 hematology analyzer to quantitatively measure hematologic traits.

#### Immunologic Traits

Peripheral blood samples were analyzed by flow cytometry using FlowJo™ v9.9.6 Software (BD Life Sciences) to determine the frequency of major circulating immune cell subsets. Analysis was performed before the start of dietary interventions at 5 months, then at 16 and 24 months of age.

#### Glucose

At the flow cytometry blood collections, mice were fasted for 4 h and glucose was measured using the OneTouch Ultra2 glucose meter from LifeScan along with OneTouch Ultra test strips.

#### Lifespan

Lifespan protocols for the Collaborative Cross Longitudinal Study were the same as those for the DO Longitudinal Study [6]. Mice were routinely evaluated for pre-specified moribund criteria. If necessary, preemptive euthanasia was performed to prevent suffering; mice euthanized or found dead were classified as deaths in survival analysis.

### Statistical analysis

Initial data quality control included identifying and resolving equipment miscalibration, mislabeled animals, and technically impossible values. Quantitative assays including body weight and temperature were explored for outliers. If we could not manually correct data points using laboratory records, they were treated as missing. Mice that did not survive sample collection (n=2) and/or to intervention onset at 6 months (n=32) were excluded from analyses, 21 males (*n_AL_*=12, *n_IF_* =9) and 12 females (*n_AL_*=6, *n_IF_* =6). The study was powered to detect global strain effects across the 10 inbred strains, but not pairwise differences in specific strains. Results from exploratory post hoc stratified analyses within strain should be considered preliminary. Analyses were conducted using R verson 4.3.3 [39].

#### Lifespan

We computed descriptive statistics to summarize lifespan data for each of the 10 Collaborative Cross inbred strains, stratified by sex and diet. Strain-specific Kaplan-Meier curves and pairwise log-rank tests estimated strain effects on lifespan. Cox proportional hazards regression models within each strain estimated the effects of treatment group, sex, and their interaction on longevity. Cox proportional hazards regression models across strains estimated average effects of treatment group, sex, and their interaction on longevity by including strain as a stratification term. Sensitivity analyses included a random intercept term for housing identifier to account for intra-cage correlations. Kaplan-Meier curves by treatment group and sex, along with Wald tests, were annotated with p-values for diet, sex, and diet-by-sex interaction effects. Non-parametric tests assessed diet effects on median lifespan within strain-sex strata. Median lifespan was estimated as the 50th percentile from survival curves using the survfit function in R’s survival package. Restricted mean survival time analysis was implemented to summarize diet effects as mean lifespan difference in months via the survRM2 R package [40, 41]. To test IF’s modulation of lifespan variability, we computed coefficients of variation (CV) for each strain by treatment group and conducted a Wilcox signed rank test to detect differences in CV by treatment group.

#### Heritability

Genetic variation in response to dietary intervention is likely attributable to large numbers of loci with small effects. For the genetic analysis of lifespan, we assessed heritability, i.e., the proportion of outcome variation in a population explained by genetic relationships. Heritability can be decomposed into multiple components, such as the proportion of variation explained by all genetic effects (broad-sense heritability, or *H*^2^) and the proportion of variation explained by additive genetic effects (narrow-sense heritability, or *h*^2^) [42]. For studies of inbred strains with replicates, an intraclass correlation is an appropriate estimator of heritability, e.g., [43]. As demonstrated in [44], replicates should not be reduced to strain-level summaries for the purpose of estimating heritability due to upward bias.

Heritability estimates for lifespan in the CC mouse population were obtained using linear mixed-effects models implemented via the lme4qtl package in R, which extends lme4 to accommodate custom covariance structures (v.0.2.2) [45]. Broad-sense heritability (*H*^2^) was estimated using a random intercept model of the form:

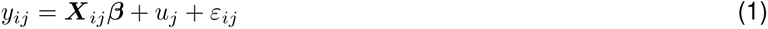

where *y_ij_* is the survival time for replicate i in strain j, ***X****_ij_* includes fixed effects for sex, diet, and their interaction, 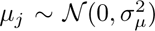 captures strain-specific random effects, and *ε_ij_* ∼ N (0*, σ*^2^) is residual error. Narrow-sense heritability (*h*^2^) was estimated by incorporating a strain-level additive genetic relationship matrix **K**, derived from CC founder haplotype probabilities using the qtl2 package. A subset of the matrix (for 10 strains included in the current study) was rescaled to ensure mean diagonal equaled 1. The model was fit as:

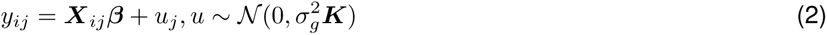

Point estimate of narrow-sense heritability was extracted using the VarProp() function, and confidence intervals were computed via likelihood profiling as implemented in lme4, which is more robust than Wald-type intervals based on standard errors where estimates are bounded, as is the case for the lme4 implementation of variance components.

## Mixed effect models

### Modeling strategy

We fit linear mixed models between outcome and study factors using the lmer function from the lme4 package in R with default control parameters. The modeling strategy can be summarized as follows:

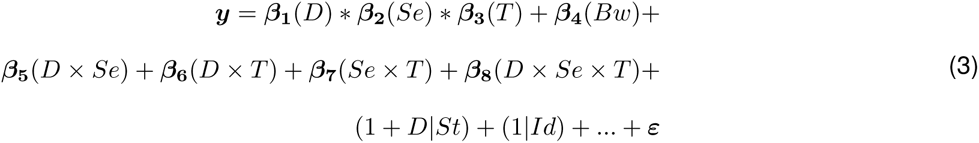

where ***y*** is rank normal transformed outcome, *D* is dietary assignment (2-day IF/ad lib), *Se* is sex (male/female), *T* is scheduled timepoint, *Bw* is body weight at the date preceding and closest to 6 months (intervention start), *St* is strain, and *Id* is a mouse-specific identifier. Batch effects not specified above (…) included additive random intercept terms for collection date (all phenotypes) and coat color and experimenter (frailty). Model complexity reflects the highly factorial nature of the study design and accounts for potential diet-by-sex-by-timepoint interaction, confounding by initial pre-treatment body weight, gene-by-treatment interaction, clustered error variance due to replicated measurement, and batch effects.

### Quantification of trajectories

For some outcomes it was possible summarize y at the individual mouse level. Bodyweight trajectories were summarized per mouse as mean mass (MM) and total AUC (+AUC); +AUC was computed with the trapezoidal rule via the integrate.xy() function of the sfsmisc R package (v.1.1-16). Per-mouse frailty trajectories were summarized as linear coefficients (slopes) on PLL scale to describe pace of aging, and final FI score was used as a proxy for lifetime multisystem frailty accumulation. Per-mouse temperature trajectories were likewise summarized as linear coefficients (slopes) on proportion of life lived (PLL [scaled age]). For these phenotypes, a simplified model was specified without fixed effects for timepoint, random effects for Id, or random effects for collection date (batch).

### Estimation of population-averaged intervention effect

For the models described above, corresponding p-values for diet fixed effect overall and within sex (and timepoint, where modeled) were computed via contrasts of model-based means estimated via the emmeans package in R. Whereas stratifying can lead to loss of information and reduced statistical power, applying pairwise tests within a single model framework provides more accurate and reliable comparisons by accounting for the overall data structure. To control for the false discovery rate across multiple phenotypes, we applied the Benjamini-Hochberg (BH) procedure to adjust diet p-values.

### Estimation of GxT from random effects

To estimate how genetic background interacts with dietary treatment (GxT interaction), we tested for diet effect heterogeneity across strains via the ranova function from the lmerTest package in R. Correlation between random effect for strain and random diet effect within strain (IF v AL), denoted as “*ρ*”, indicated how strain-specific diet effect varied across strain-specific estimates for baseline control. To control for the false discovery rate, we applied the Benjamini-Hochberg (BH) procedure to adjust p-values for diet random effect and, separately, *ρ* across multiple phenotypes.

### Mouse model selection

We applied random effects post-estimation to the panel of mouse strains as models for heterogeneous responses to intermittent fasting. Best linear unbiased predictions (BLUPs) provide empirical Bayes (EB) estimates of random effects, *ζ_EB_*, offering a principled way to quantify strain-level deviations in IF response from population mean response while accounting for uncertainty and shrinkage toward the overall mean. In this context, *ζ_EB_* capture the magnitude and direction of each strain’s response to intermittent fasting (IF) conditional on the fitted model structure. To generate interpretable strain-specific diet effect scores, we summed *ζ_EB_* and population mean diet response computed from post-estimated model contrasts taking sex-by-diet interaction into account. For hematologic and immunologic phenotypes, total strain-specific diet effects scaled within strain were grouped into clusters via unsupervised learning and co-displayed with heatmaps organized by health outcome and strain generated via the tidyHeatmap package in R, revealing groups of strains with distinctive treatment response.

#### Comparative analysis

In a comparative analysis of IF effects across genetically diverse mouse populations (CC and DO), we conducted parallel analyses of data from the outbred DRiDO study and the inbred CC panel. We reanalyzed a subset of the DRiDO dataset, restricting to outbred mice assigned to either ad libitum (AL) or 2-day intermittent fasting (IF) protocols and applied our phenotypic response analysis pipeline to this subset to obtain comparable statistical estimates of IF effects. Thus in the current manuscript “IF” refers to the 2-day protocol unless otherwise specified. The analysis pipeline is as described for the CC study in Methods: Modeling strategy, except that batch, sex, and strain were excluded (DRiDO mice were outbred females, data were pre-adjusted for batch). Results were compared to female-specific contrasts post-estimated from linear mixed models (LMMs) in the CC dataset, as described in Methods: Estimation of population-averaged intervention effect.

Reanalysis focused on a shared subset of trait domains: body weight, body composition, frailty, hematology, and immunology. Hematologic and immunologic traits were selected based on their prominence in the original DRiDO publication. Comparability across the two studies for body composition analysis and immunophenotyping is lesser than for other phenotypic domains as scientific staff implemented new assessment modalities in the period between conducting DRiDO and the CC, transitioning from dual-energy x-ray absorptiometry (DEXA) to nuclear magnetic resonance body composition assessment EchoMRI-3-in-1 Whole Body Resonance Analyzer and from fluorescence-activated cell sorting (FACS) to flow cytometry-based immunophenotyping.

#### Non-linear models of body temperature

To investigate the interaction between sex, diet, and age on phenotypes, we fit linear mixed models (LMMs) with an ordinal timepoint variable (e.g., timepoint ∈ 0, 1) to represent the study’s discrete measurement intervals [see Methods: Modeling strategy], or we summarized the trait across the lifespan (e.g., slope of frailty score) [see: Methods: Quantification of trajectories]. For temperature, which was collected with greater temporal density than other phenotypes (5 timepoints), we reformulated time as a continuous variable by normalizing age at sampling relative to lifespan and applied generalized additive mixed model (GAMM) methods.

Prior to rescaling age, the data structure had fewer unique time intervals and a more constrained fixed-effects space, i.e., was more amenable to reliable estimation of complex random effects. In contrast, the continuous-time model, while richer in biological insight, imposes greater demands on model identifiability and stability. We omitted the random slope for diet, reflecting a pragmatic balance between model complexity, data limitations, and the interpretive clarity of the results. Formal statistical tests for random effects in GAMMs remain underdeveloped, thus we also re-specified strain as a fixed (vs. random) effect to enable hypothesis testing of strain and age-by-strain interactions. I.e., in transitioning to a continuous-time framework, we simplified the random effect model structure to include only a random intercept for mouseID.

GAMMs were fit using the mgcv R package (v.1.9-1) [46]. Random intercepts for were implemented via a smooth term with basis type ‘re’, allowing for replicate-specific deviations. Approximate significance of smooth terms was assessed using a frequentist approximation based on F-tests. Tests are approximate in that they rely on large sample assumptions and do not fully account for the uncertainty introduced by the smoothing parameter selection process, i.e., the data-driven selection of effective degrees of freedom.

#### Longevity biomarkers

To identify traits that are associated with lifespan, we performed regression analysis on lifespan with traits at each timepoint after adjusting for effects of diet, sex, strain, batch, and body weight. Weekly body weight outcome was modeled as MM and +AUC (defined above) rather than by timepoint. All continuous variables were rank normalized before model fitting, resulting in standardized beta coefficients. We performed likelihood ratio tests to obtain p-values for the adjusted associations via the anova function from the lmerTest R package (v.3.1-3) using Satterthwaite’s method for computing denominator degrees of freedom and F-statistics. We applied an FDR adjustment to each test across traits and timepoints (one-step Benjamini–Hochberg method).

## Acknowledgments

This work was supported by the National Institute on Aging award number P30 AG038070 to Gary Churchill and Ron Korstanje (co-PI). DRiDO data used in this study were previously collected with support from Calico Life Sciences LLC (Dietary Intervention of Aging in Genetically Diverse Mice, sponsored research funding number CALICO-GAC-06), and are reused here with permission. We thank Andrew Deighan for providing assistance with data curation. We acknowledge the JAX Nathan Shock Center Animal and Phenotyping Core team for their expertise in animal husbandry, data collection, and data curation, the JAX Center for Biometric Analysis for overseeing NMR data collection and processing, and the JAX Flow Cytometry Service for assistance with flow cytometry panel design, data acquisition, and gating analysis. The JAX Flow Cytometry Service is a Shared Resource of the JAX Cancer Center (P30 CA034196).

## Author Contributions

AL: methodology, formal analysis, writing. LR: project administration, data curation. WS: methodology, resources, writing. APW: formal analysis, writing. RK: funding acquisition, project administration, writing. GC: conceptualization, funding acquisition, data acquisition, project administration, methodology, formal analysis, writing.

## Data Availability Statement

All analyses were performed using the R statistical programming language [39]. Data and code used to generate tables, figures, and reported results can be found on Figshare (DOI:10.6084/m9.figshare.30013813). CC strain genotypes were obtained from https://www.jax.org/research-and-faculty/genetic-diversity-initiative/tools-data/diversity-outbred-reference-data. Data for DRiDO phenotypes can be found on the online data repository for the anchor manuscript [47].

