## Supporting Information for "Genetic regulation of fasting-induced longevity effects"

**Supplementary Figure 1: Sex-specific effects of dietary intervention on lifespan across genetically diverse mouse strains.** A) Kaplan-Meier curves and interaction contrasts for lifespan response to diet among males (dashed) and females (solid) reveal inter-strain variation in IF response. Abbreviations: mo. = months, int = interaction. B) Data from the CC Longitudinal Study show sexual dimorphism and genetic effects in IF response. Visualization highlights the interaction between sex, diet, and genetic background in modulating longevity. Lifespan is plotted for male and female mice, grouped by dietary regimen and genetic strain. Data are shown for ad libitum feeding (○) and 2-day intermittent fasting (Δ) and plotted by sex as median ± IQR.

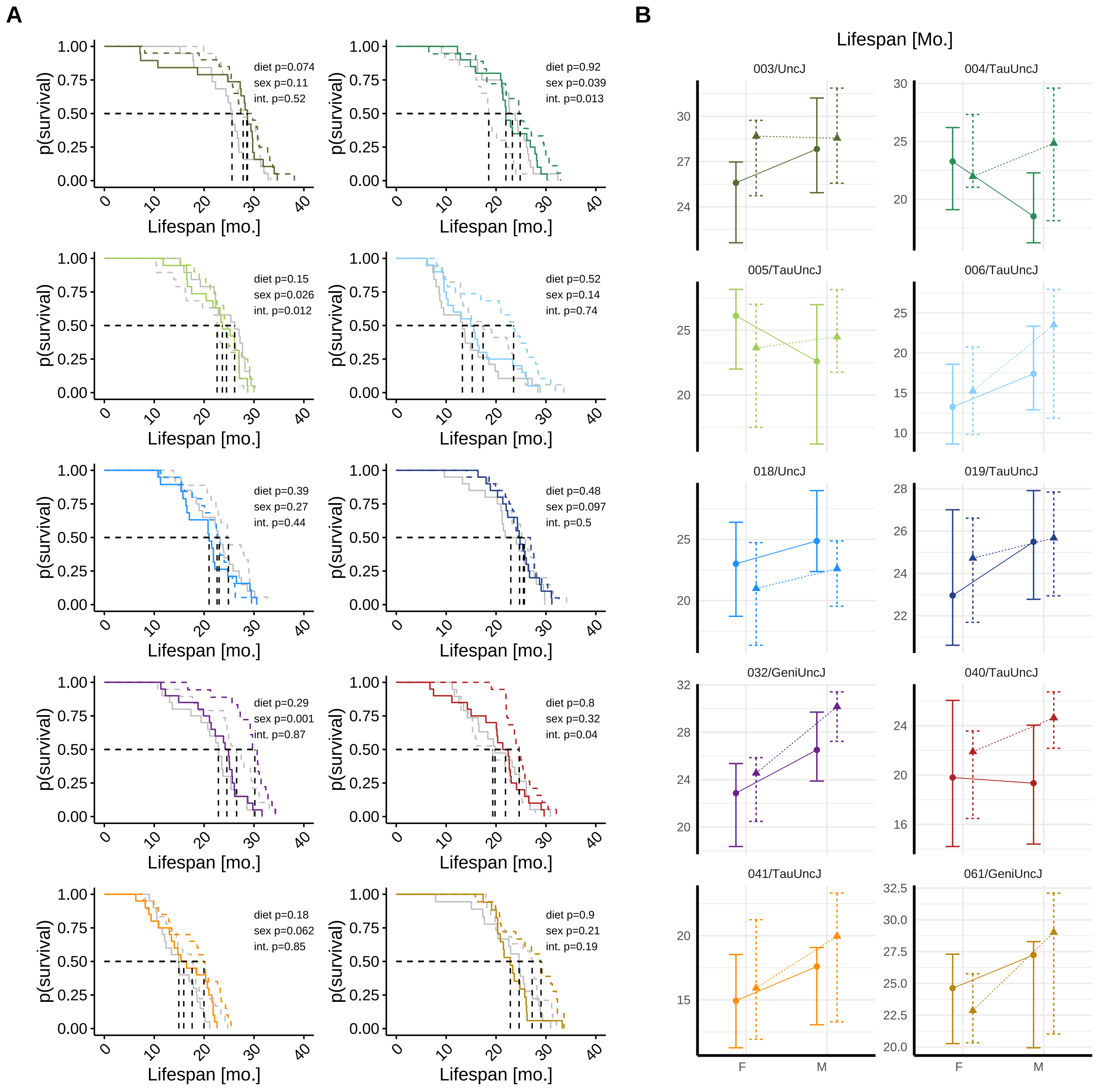

**Supplementary Figure 2: Follow-up by physiologic trait, year, sex, and intervention group.** For phenotyping domains assessed multiple times per year, e.g., frailty, any measurement within the study year was counted as continued follow-up. In total, longitudinal phenotyping included over 66,000 body weight measurements, 2,000 frailty assessments, and 1,000 each of hematology, immunology, metabolic phenotype, and glucose tolerance assays. Abbreviations: Fasted.Gl=fasted glucose, AL=*ad libitum*, IF=intermittent fasting.

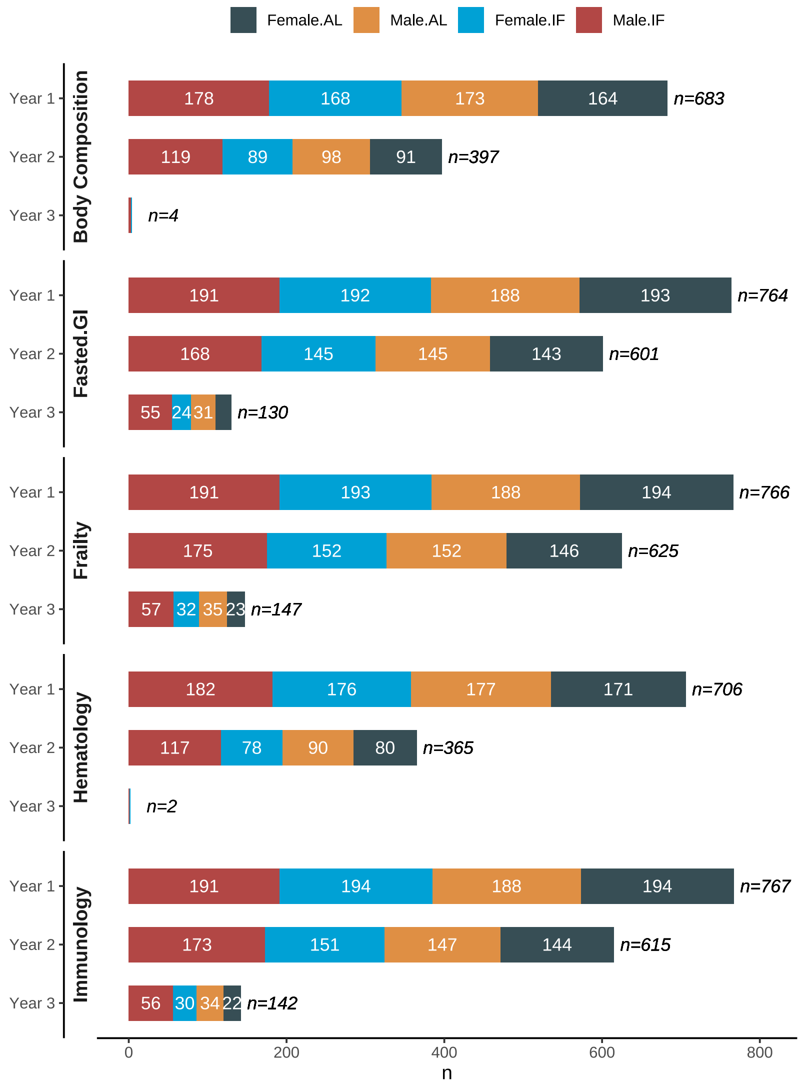

**Supplementary Figure 3: Metabolic phenotype response to intermittent fasting (IF) is influenced by genetic background.** Nuclear magnetic resonance (NMR) body composition assays were conducted at 10 and 22 months (n>1k). Month 22 NMR data were plotted as mean ± SE adiposity [%] (A) and lean mass [g] (B) for male and female mice, grouped by dietary regimen (IF [Δ] or AL [○]) and genetic strain. Early lean mass loss in response to IF was observed with varied risk across strains and sexes. Some strains (e.g., 005/TauUncJ, 006/TauUncJ) showed minimal total mass change but significant lean mass loss, underscoring the importance of composition-specific metrics for characterizing metabolic phenotype. Plots for 10 month timepoint shown in **Fig. 3a-b**.

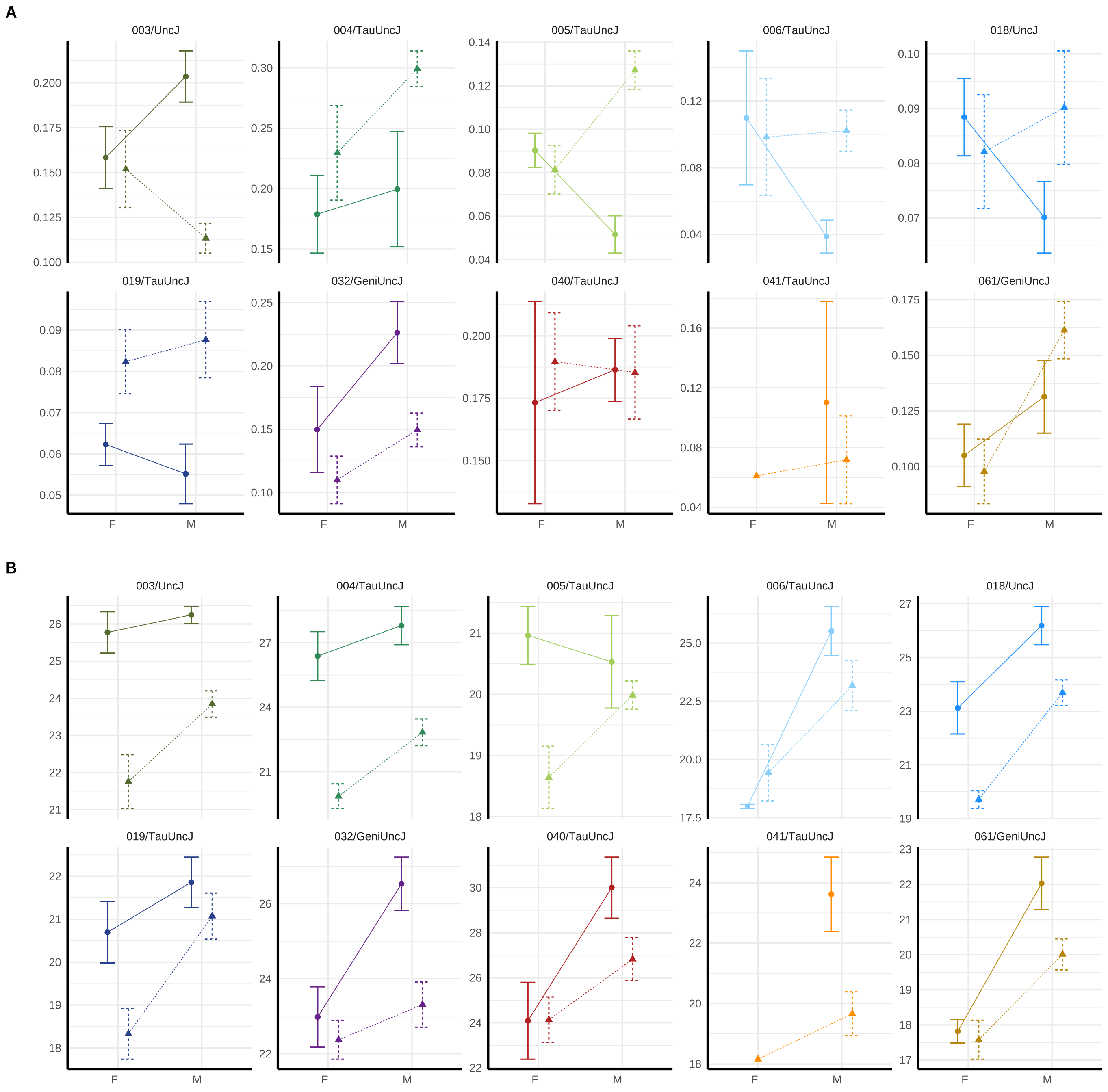

**Supplementary Figure 4: Lifetime incidence of individual health deficits highlights several frailty indicators characterized by low susceptibility and minimal influence from genetic or dietary factors.** Twenty-seven serially collected non-invasive biomarkers of frailty were measured at weeks 21 (preintervention), 43 (intervention onset), 95, 121, and 147 weeks (n>2,000). The comparison of cumulative incidence of individual health deficits across intervention groups (IF = color, AL = gray) reveals several physiologic systems with low aggregate risk of frailty accumulation across the lifespan regardless of genetic background or exposure to IF dietary regimen. Note: Remaining items shown in **Fig. 4**. Frailty index items were binarized as severe/nonsevere.

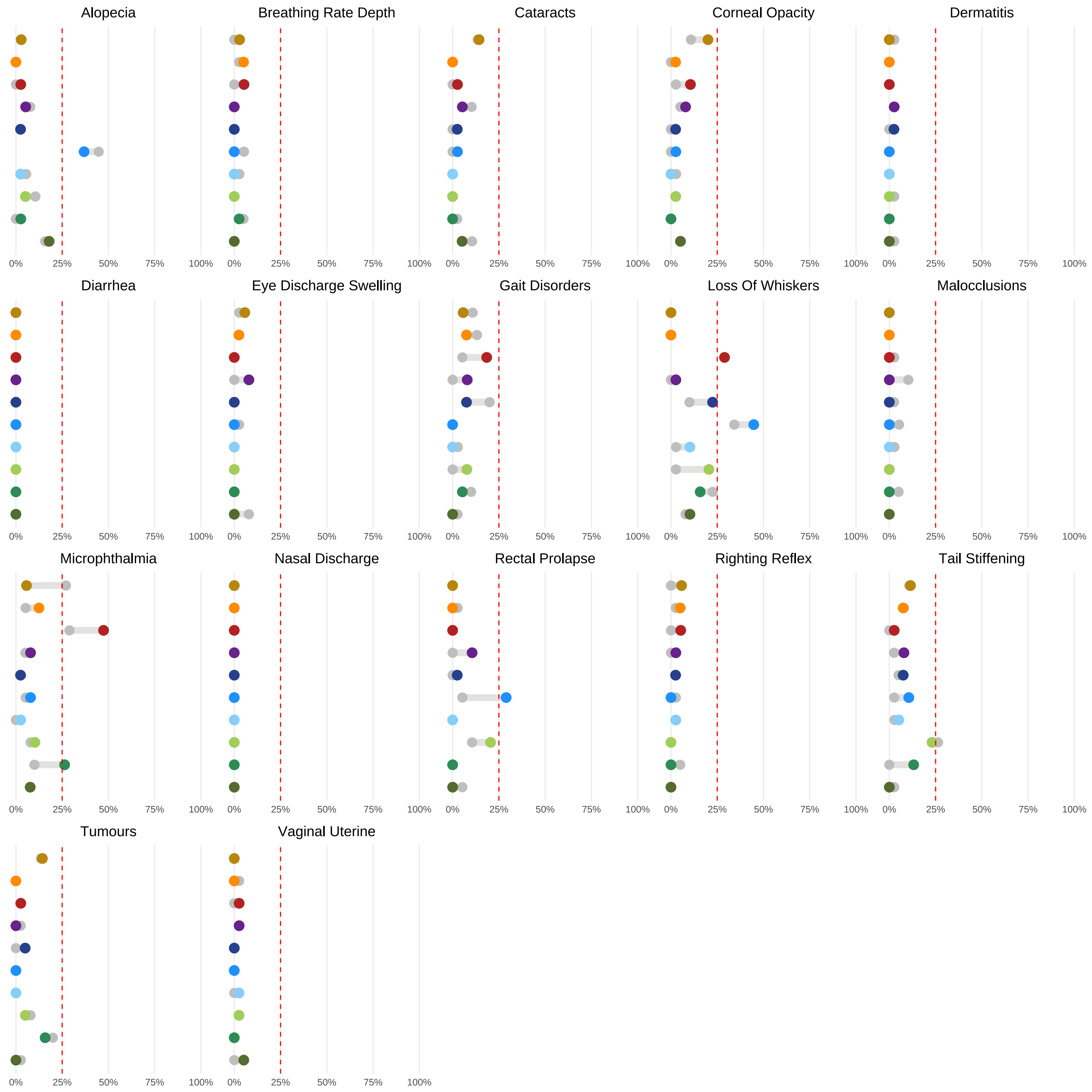

**Supplementary Figure 5: Genetic signatures of hematologic response to IF in late life.** Year 2 red blood cell distribution width (RDW CV) data (A) and mean corpuscular volume (MCV) (B) were plotted as mean $\pm$ SE for male and female mice grouped by dietary regimen (IF [$\Delta$] or AL [$\circ$]) and genetic strain, demonstrating sexually dimorphic IF response at 22 months. Note: 10-month data for RDW and MCV are shown in **Fig. 5**.

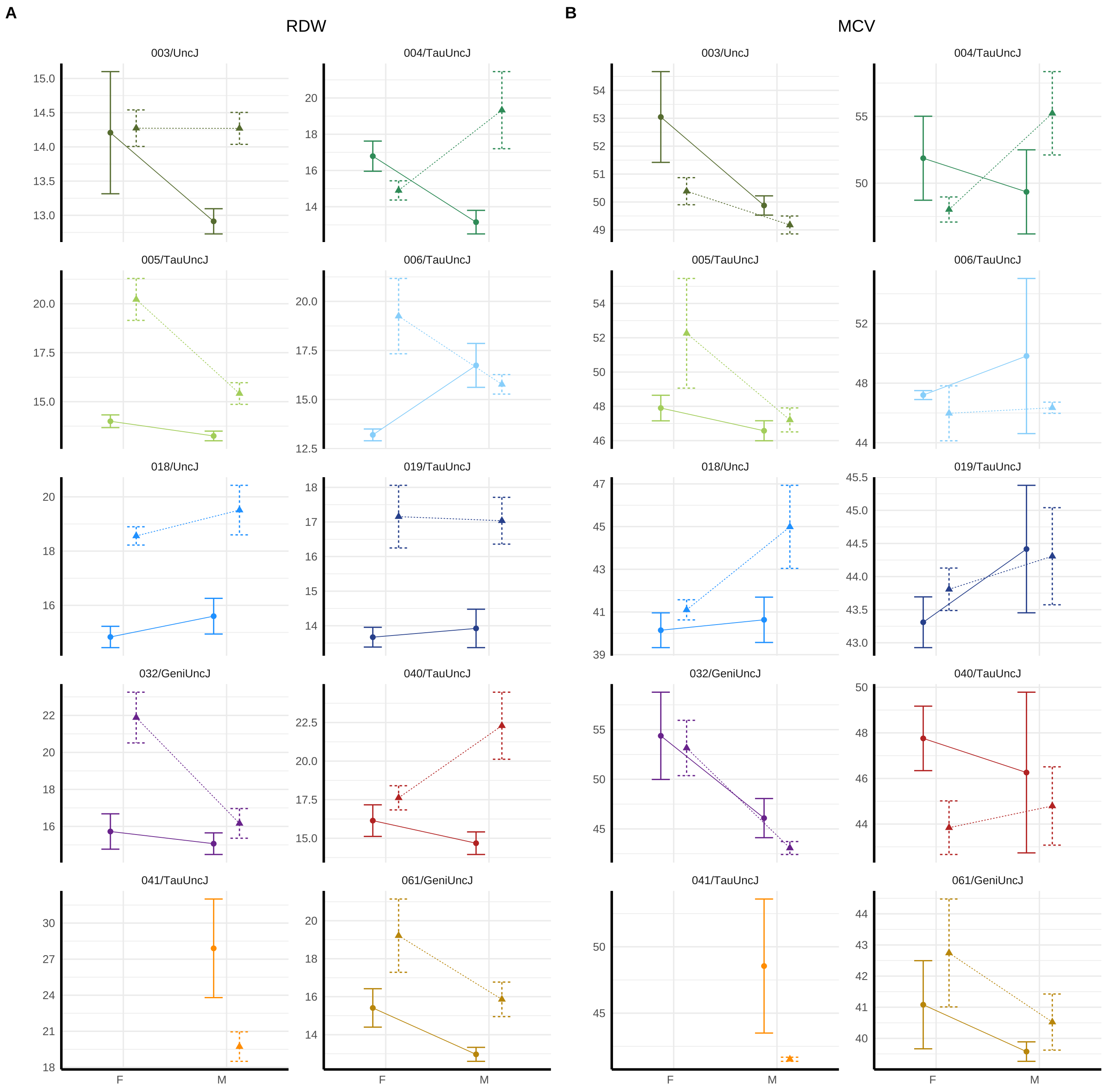

**Supplementary Figure 6: Varying hematologic IF responses clustered by strain demonstrate how deep longitudinal phenotyping in the Collaborative Cross enables discovery of models for intervention response.** Heatmaps compare diet effects by hematologic outcome for males and females at year 1 and year 2. Clustering partitions and dendrograms reveal latent groups of strains with similarly patterned hematologic intervention response. Notes: Row (hematologic trait) order for 45-week (year 1) female heatmap [top left; see row-wise dendrogram] determined by cluster analysis; row order for remaining plots were pre-specified without clustering to allow for visual comparison across study strata. Column (strain) order determined by clustering analysis for each heatmap separately with partitions to group strains with similar hematologic profile response to IF. The reference group for strain-specific diet effects shown in the heatmap is parameterized such that the diet comparison is IF-AL. Negative values indicate adjusted mean outcome for IF is lower than for the reference level AL. Abbreviations for hematologic traits are defined in Supplementary Table S6.

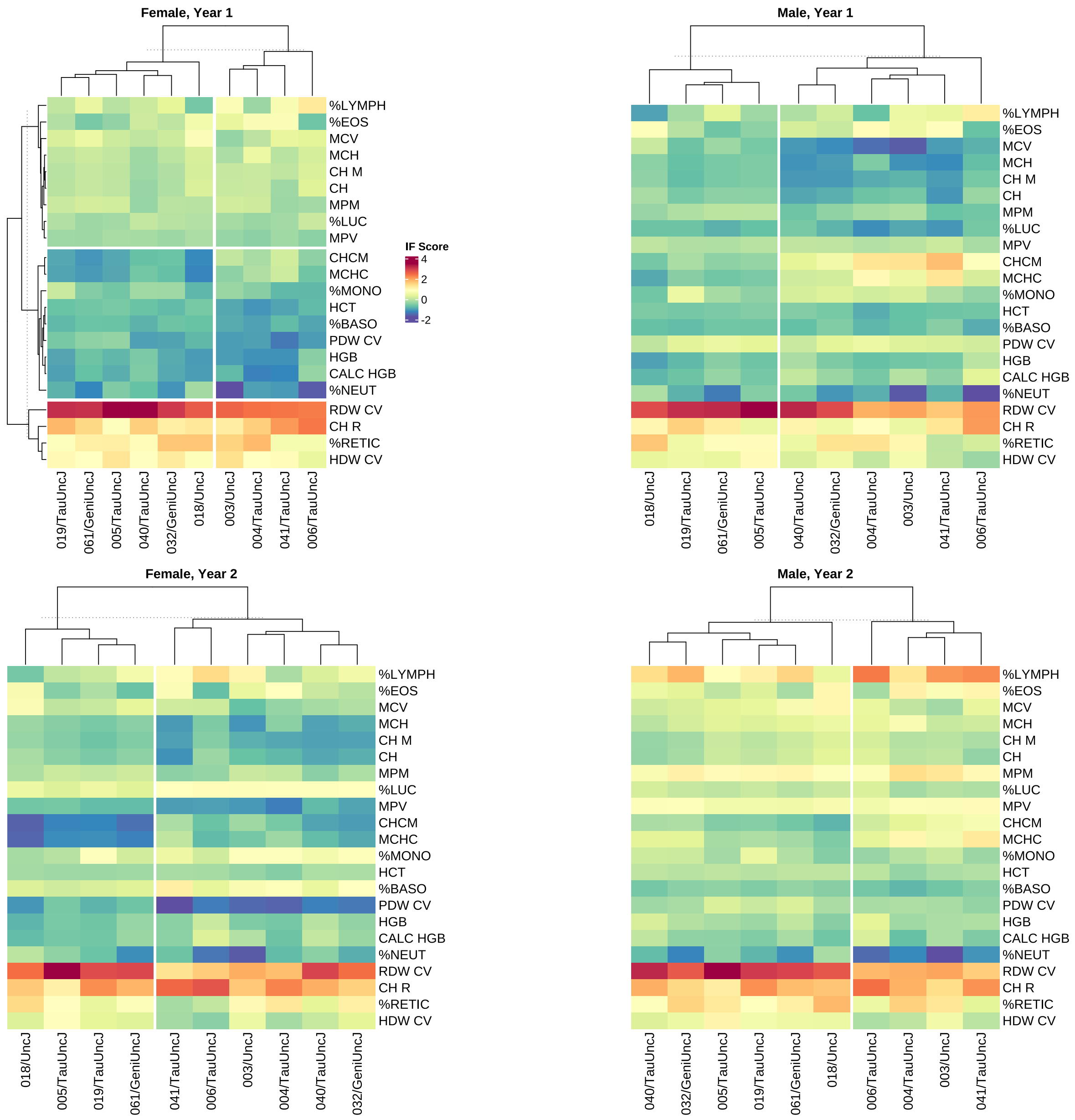

**Supplementary Figure 7: Varying immunologic IF responses clustered by strain demonstrate how deep longitudinal phenotyping in the Collaborative Cross enables discovery of models for intervention response.** Heatmaps compare diet effects by immunologic outcome for males and females at year 1 and year 2. Clustering partitions and dendrograms reveal latent groups of strains with similarly patterned immunologic intervention response. Notes: Row (immunologic trait) order for 45-week (year 1) female heatmap [top left; see row-wise dendrogram] determined by cluster analysis; row order for remaining plots were pre-specified without clustering to allow for visual comparison across study strata. Column (strain) order determined by clustering analysis for each heatmap separately with partitions to group strains with similar immunologic profile response to IF. The reference group for strain-specific diet effects shown in the heatmap is parameterized such that the diet comparison is IF-AL. Negative values indicate adjusted mean outcome for IF is lower than for the reference level AL. Abbreviations for immunologic traits are defined in Supplementary Table S6.

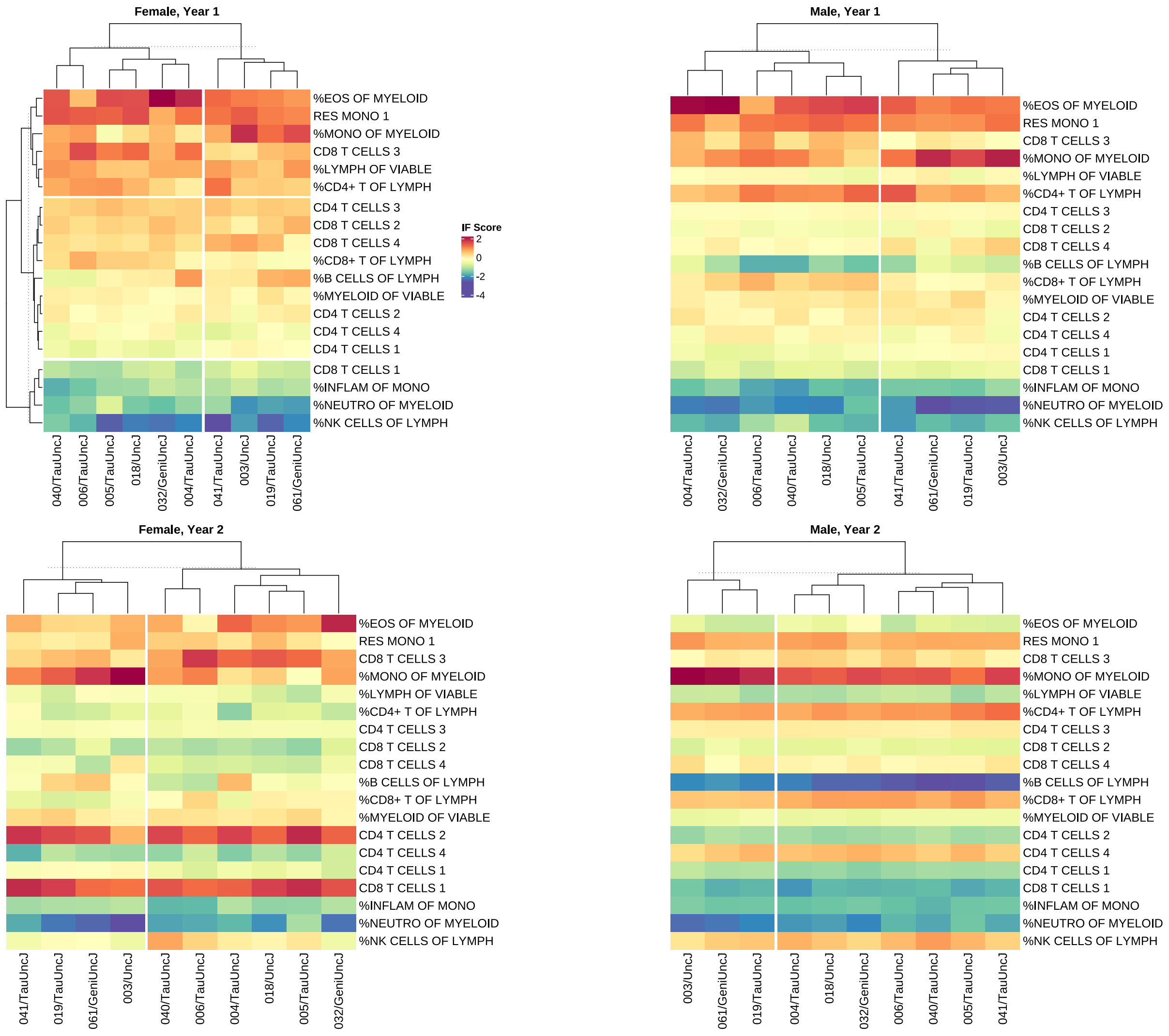

**Supplementary Table 1:** Comparison of diet effects on lifespan by sex.

**(a)** Median lifespan by diet and sex. Significance (p-value) of comparisons (1df) between diet groups from $\chi^2$ tests.

| | IF | AL | IF-AL | p |
|---|---|---|---|---|
| Female | 21.97 | 22.24 | -0.26 | 0.92 |
| Males | 24.84 | 23.17 | 1.66 | 0.02 |

**(b)** Maximum lifespan (90% survival) by diet and sex. Significance (p-value) of comparisons (1df) between diet groups from $\chi^2$ tests.

| | IF | AL | IF-AL | p |
|---|---|---|---|---|
| Females | 29.05 | 28.59 | 0.46 | 0.49 |
| Males | 31.41 | 30.79 | 0.62 | 0.30 |

**(c)** Mean lifespan by diet and sex. Significance (p-value) of comparisons (1df) between diet groups from Restricted Mean Survival Time (RMST) analysis.

| | IF | AL | IF-AL (95% CI) | p |
|---|---|---|---|---|
| Females | 21.34 | 21.01 | 0.334 (-0.94, 1.61) | 0.607 |
| Males | 24.37 | 22.35 | 2.02 (0.76, 3.28) | 0.002 |

**Supplementary Table 2:** Descriptive statistics for lifespan. Survival time for specified quantiles were determined using the Kaplan-Meier estimator, with the desired percentile extracted via the quantile function applied to the survival curve. Maximum lifespan defined as 90th percentile lifespan within study strata. Values are rounded to the nearest 10th of a month.

**(a)** Lifespan summarized by strain and sex.

| Strain | p50_F | p50_M | p25_F | p25_M | p75_F | p75_M | max_F | max_M |
|---|---|---|---|---|---|---|---|---|
| 003/UncJ | 26.9 | 27.8 | 22.3 | 25.4 | 29.6 | 31.2 | 32.0 | 33.3 |
| 004/TauUncJ | 22.6 | 21.0 | 21.0 | 17.4 | 26.4 | 25.0 | 28.2 | 30.6 |
| 005/TauUncJ | 25.3 | 23.9 | 20.4 | 19.7 | 27.2 | 27.0 | 28.8 | 29.1 |
| 006/TauUncJ | 13.8 | 21.6 | 9.4 | 12.9 | 18.6 | 25.9 | 26.4 | 28.5 |
| 018/UncJ | 21.9 | 23.4 | 17.0 | 21.2 | 25.8 | 26.2 | 29.4 | 29.3 |
| 019/TauUncJ | 24.5 | 25.5 | 21.1 | 22.8 | 26.6 | 27.8 | 29.6 | 31.0 |
| 032/GeniUncJ | 23.5 | 29.2 | 19.6 | 24.9 | 25.6 | 31.0 | 28.7 | 33.1 |
| 040/TauUncJ | 21.3 | 22.6 | 15.0 | 19.1 | 24.7 | 25.3 | 26.7 | 28.2 |
| 041/TauUncJ | 15.0 | 18.5 | 11.3 | 13.1 | 20.0 | 22.7 | 21.7 | 24.5 |
| 061/GeniUncJ | 23.4 | 27.2 | 20.3 | 21.0 | 26.0 | 31.2 | 28.8 | 32.3 |

**(b)** Lifespan summarized by strain, sex, and diet.

| Strain | Diet | p50_F | p50_M | p25_F | p25_M | p75_F | p75_M | max_F | max_M |
|---|---|---|---|---|---|---|---|---|---|
| 003/UncJ | AL | 25.6 | 27.8 | 21.6 | 24.9 | 27.0 | 31.2 | 32.0 | 32.3 |
| 003/UncJ | IF | 28.7 | 28.6 | 24.7 | 25.6 | 29.7 | 31.9 | 34.0 | 33.9 |
| 004/TauUncJ | AL | 23.3 | 18.5 | 19.1 | 16.2 | 26.2 | 22.3 | 27.2 | 23.7 |
| 004/TauUncJ | IF | 22.0 | 24.8 | 21.0 | 18.2 | 27.3 | 29.6 | 28.6 | 32.3 |
| 005/TauUncJ | AL | 26.1 | 22.6 | 22.0 | 16.2 | 28.2 | 27.0 | 29.4 | 27.6 |
| 005/TauUncJ | IF | 23.7 | 24.5 | 17.5 | 21.8 | 27.0 | 28.1 | 28.7 | 29.6 |
| 006/TauUncJ | AL | 13.3 | 17.4 | 8.6 | 12.9 | 18.6 | 23.4 | 27.2 | 25.9 |
| 006/TauUncJ | IF | 15.2 | 23.5 | 9.8 | 11.8 | 20.8 | 28.0 | 26.2 | 31.0 |
| 018/UncJ | AL | 23.0 | 24.9 | 18.7 | 22.4 | 26.4 | 29.0 | 29.4 | 29.6 |
| 018/UncJ | IF | 21.0 | 22.6 | 16.3 | 19.5 | 24.7 | 24.9 | 29.4 | 26.2 |
| 019/TauUncJ | AL | 23.0 | 25.5 | 20.6 | 22.8 | 27.0 | 27.9 | 29.6 | 31.1 |
| 019/TauUncJ | IF | 24.7 | 25.7 | 21.7 | 22.9 | 26.6 | 27.8 | 30.1 | 30.8 |
| 032/GeniUncJ | AL | 22.9 | 26.5 | 18.4 | 23.9 | 25.4 | 29.7 | 28.6 | 33.1 |
| 032/GeniUncJ | IF | 24.6 | 30.2 | 20.5 | 27.2 | 25.9 | 31.4 | 29.3 | 33.6 |
| 040/TauUncJ | AL | 19.8 | 19.3 | 14.2 | 14.4 | 26.1 | 24.0 | 26.7 | 26.8 |
| 040/TauUncJ | IF | 21.9 | 24.6 | 16.5 | 22.2 | 23.6 | 26.7 | 27.8 | 30.4 |
| 041/TauUncJ | AL | 14.9 | 17.6 | 11.3 | 13.1 | 18.5 | 19.1 | 20.0 | 24.2 |
| 041/TauUncJ | IF | 15.9 | 20.0 | 11.9 | 13.3 | 21.2 | 23.3 | 22.0 | 24.7 |
| 061/GeniUncJ | AL | 24.6 | 27.2 | 20.3 | 19.9 | 27.3 | 28.3 | 29.3 | 31.4 |
| 061/GeniUncJ | IF | 22.9 | 29.0 | 20.3 | 21.0 | 25.8 | 32.1 | 26.2 | 33.6 |

**Supplemental Table 3: Pairwise comparison of strain effects on lifespan.**
Significance (p-value) of all pairwise comparisons (1df) between strain groups from log rank tests. Overall (9df) significance p<2.2e-16.

| | 003/ UncJ | 004/ TauUncJ | 005/ TauUncJ | 006/ TauUncJ | 018/ UncJ | 019/ TauUncJ | 032/ GeniUncJ | 040/ TauUncJ | 041/ TauUncJ |
|---|---|---|---|---|---|---|---|---|---|
| 004/TauUncJ | 2.8e-07 | | | | | | | | |
| 005/TauUncJ | 2.1e-07 | 0.7686 | | | | | | | |
| 006/TauUncJ | 2.5e-11 | 0.0167 | 0.0026 | | | | | | |
| 018/UncJ | 9.2e-08 | 0.9646 | 0.9323 | 0.0087 | | | | | |
| 019/TauUncJ | 0.00055 | 0.0486 | 0.0802 | 2.7e-05 | 0.0466 | | | | |
| 032/GeniUncJ | 0.04860 | 0.0022 | 0.0022 | 2.8e-07 | 0.0011 | 0.1588 | | | |
| 040/TauUncJ | 1.3e-10 | 0.2886 | 0.0957 | 0.1159 | 0.2707 | 0.0016 | 2.8e-05 | | |
| 041/TauUncJ | <2e-16 | 2.8e-09 | 5.6e-15 | 0.0319 | 3.2e-11 | <2e-16 | <2e-16 | 2.9e-08 | |
| 061/GeniUncJ | 0.02113 | 0.0085 | 0.0125 | 1.4e-06 | 0.0085 | 0.3662 | 0.69 | 0.00012 | <2e-16 |

**Supplemental Table 4: Diet effects on mean lifespan by sex and strain**

| Strain | Sex | Ratio [IF/AL] (95% CI) | p |
|---|---|---|---|
| 003/UncJ | Female | 0.55 (0.28, 1.06) | 0.074 |
| 003/UncJ | Male | 0.74 (0.39, 1.42) | 0.367 |
| 004/TauUncJ | Female | 1.03 (0.55, 1.94) | 0.915 |
| 004/TauUncJ | Male | 0.32 (0.16, 0.63) | 9e-04 |
| 005/TauUncJ | Female | 1.62 (0.84, 3.12) | 0.149 |
| 005/TauUncJ | Male | 0.46 (0.24, 0.91) | 0.026 |
| 006/TauUncJ | Female | 0.81 (0.43, 1.53) | 0.518 |
| 006/TauUncJ | Male | 0.69 (0.35, 1.36) | 0.284 |
| 018/UncJ | Female | 1.32 (0.7, 2.49) | 0.390 |
| 018/UncJ | Male | 1.9 (0.97, 3.72) | 0.059 |
| 019/TauUncJ | Female | 0.8 (0.43, 1.5) | 0.485 |
| 019/TauUncJ | Male | 1.09 (0.58, 2.05) | 0.788 |
| 032/GeniUncJ | Female | 0.71 (0.38, 1.34) | 0.294 |
| 032/GeniUncJ | Male | 0.66 (0.34, 1.28) | 0.222 |
| 040/TauUncJ | Female | 1.09 (0.58, 2.05) | 0.797 |
| 040/TauUncJ | Male | 0.41 (0.21, 0.8) | 0.009 |
| 041/TauUncJ | Female | 0.65 (0.34, 1.23) | 0.183 |
| 041/TauUncJ | Male | 0.59 (0.31, 1.15) | 0.122 |
| 061/GeniUncJ | Female | 1.04 (0.53, 2.07) | 0.900 |
| 061/GeniUncJ | Male | 0.56 (0.28, 1.11) | 0.095 |

**Supplementary Table 5: Coefficient of variation (CV) in lifespan by diet, strain, and sex**

| Strain | Diet | CV_Female | CV_Male |
|---|---|---|---|
| 003/UncJ | AL | 0.19 | 0.13 |
| 004/TauUncJ | AL | 0.25 | 0.27 |
| 005/TauUncJ | AL | 0.19 | 0.28 |
| 006/TauUncJ | AL | 0.47 | 0.40 |
| 018/UncJ | AL | 0.24 | 0.21 |
| 019/TauUncJ | AL | 0.25 | 0.18 |
| 032/GeniUncJ | AL | 0.26 | 0.24 |
| 040/TauUncJ | AL | 0.29 | 0.28 |
| 041/TauUncJ | AL | 0.26 | 0.30 |
| 061/GeniUncJ | AL | 0.25 | 0.19 |
| 003/UncJ | IF | 0.33 | 0.23 |
| 004/TauUncJ | IF | 0.24 | 0.28 |
| 005/TauUncJ | IF | 0.22 | 0.17 |
| 006/TauUncJ | IF | 0.44 | 0.37 |
| 018/UncJ | IF | 0.28 | 0.20 |
| 019/TauUncJ | IF | 0.17 | 0.18 |
| 032/GeniUncJ | IF | 0.24 | 0.15 |
| 040/TauUncJ | IF | 0.32 | 0.13 |
| 041/TauUncJ | IF | 0.33 | 0.29 |
| 061/GeniUncJ | IF | 0.16 | 0.20 |

**Supplementary Table 6: Trait definitions.**
Filename: Tab_S6.csv
Content: Description of study data.
Contains the following fields:
Task Name: corresponds to 'task_name' in dataset
Task Description: defines 'task_name'
Trait Name: corresponds to 'variable' in dataset (phenotypic trait)
Trait Description: defines 'variable'

**Supplemental Table 7: Phenotyping domains with age ranges.**
Due to scheduling constraints, it was not possible to test all mice at precisely the same ages. In this table we record for each phenotyping event (indexed by phenotyping domain and timepoint [in months]) the youngest (min_age), median (median_age), and oldest (max_age) ages at which mice were tested.

| Domain | Timepoint | min_age | med_age | max_age |
|---|---|---|---|---|
| CBC | 10.00 | 9.20 | 11.40 | 12.10 |
| CBC | 22.00 | 21.40 | 23.40 | 24.20 |
| CBC | 34.00 | 33.60 | 34.70 | 35.80 |
| FLOW | 5.00 | 3.30 | 5.20 | 5.90 |
| FLOW | 16.00 | 14.30 | 16.30 | 17.60 |
| FLOW | 28.00 | 26.20 | 28.20 | 28.80 |
| Fasted.GI | 6.00 | 3.70 | 5.60 | 6.30 |
| Fasted.GI | 17.00 | 14.70 | 16.70 | 17.20 |
| Fasted.GI | 28.00 | 17.30 | 28.70 | 29.70 |
| Frailty | 5.00 | 3.70 | 5.00 | 7.40 |
| Frailty | 10.00 | 7.90 | 10.10 | 10.90 |
| Frailty | 16.00 | 14.10 | 16.10 | 16.70 |
| Frailty | 22.00 | 20.80 | 22.00 | 23.20 |
| Frailty | 28.00 | 26.10 | 28.10 | 28.80 |
| Frailty | 33.00 | 32.10 | 33.50 | 34.10 |
| NMR | 10.00 | 8.40 | 10.60 | 11.40 |
| NMR | 23.00 | 20.60 | 22.50 | 23.20 |
| NMR | 32.00 | 32.60 | 33.60 | 34.50 |

**Supplementary Table 8: Diet effect in recombinant inbred strain panel.**
Filename: Tab_S8.csv
Content: Longitudinal analysis test statistics for diet effect on traits. File contains the following fields:
Variable: [trait domain]_[trait name].
Contrast: Direction of effect estimated
Timepoint: Timepoint in weeks. Not applicable for traits summarized at the individual level described in Methods: Summarization of trajectories.
Estimate, Estimate_Male, Estimate_Female: estimate of adjusted diet association, as described in Methods: Estimation of population-averaged intervention effect. Adjustment factors as described in Methods: Modeling strategy.
p.value, p.value_Female, p.value_Male: p-value for estimates. Adjustment factors as described in Methods: Modeling strategy.
q.value, q.value_Female, q.value_Male: false discovery rate adjusted significance.

**Supplementary Table 9: Diet effect in recombinant inbred strain panel.**
Filename: Tab_S9.csv
Content: Estimation of gene-by-treatment interaction as described in Methods: “Estimation of GxT from random effects.”
Variable: [trait domain]_[trait name].
p.value_sd_diet: p-value for diet effect heterogeneity across strains.
q.value_sd_diet: false discovery rate adjusted of above.
Coefficient_[strain]: diet random effect, corresponding to the deviation from fixed effect.

**Supplementary Table 10: Strain and diet random effect correlation.**
Filename: Tab_S10.csv
Content: Estimation of correlation between strain and diet random effects as described in Methods: “Estimation of GxT from random effects.”
Variable: [trait domain]_[trait name].
cor_diet_strain_res: correlation between strain and diet random effects.

**Supplementary Table 11: Diet effect in outbred population.**

Filename: Tab_S11.csv

Content: Longitudinal analysis test statistics for diet effect on traits in outbred population. File contains the following fields:

Variable: Trait name

Contrast: Direction of effect estimated

Timepoint: Timepoint in years. Not applicable for traits summarized at the individual level described in Methods: Summarization of trajectories.

Estimate: Estimate of adjusted diet association, as described in Methods: Comparative analysis. Note female-specific, per study design. A rank-normalized effect sizes (RNES)—defined as the estimated marginal mean difference on the rank-z transformed scale.

p.value: p-value for estimates. Adjustment factors as described in Methods: Modeling strategy.

q.value: false discovery rate adjusted significance.

**Supplementary Table 12: Phenotype correlations with lifespan.**
Filename: Tab_S12.csv
Content: Lifespan association test statistics as described in Methods: “Trait Association with Lifespan.”
File contains the following fields:
Domain: Type of assay. Variable: Trait name
Timepoint: Timepoint in weeks.
p_adj: p-value of trait-lifespan association adjusted for sex, diet, strain, and body weight.
q_adj: false discovery rate adjusted of above.
b_adj: adjusted (partial) correlation of lifespan with trait.
